# Meiosis-Specific Cohesin Complexes Display Distinct and Essential Roles in Mitotic ESC Chromosomes

**DOI:** 10.1101/2021.05.03.442391

**Authors:** Eui-Hwan Choi, Young Eun Koh, Seobin Yoon, Yoonsoo Hahn, Keun P. Kim

## Abstract

Cohesin is a chromosome-associated SMC kleisin complex that mediates sister chromatid cohesion, recombination, and most chromosomal processes during mitosis and meiosis. Through high-resolution 3D-structured illumination microscopy and functional analyses, we report multiple biological processes associated with the meiosis-specific cohesin components, REC8 and STAG3, and the distinct loss of function of meiotic cohesin during the cell cycle of embryonic stem cells (ESCs). First, we show that REC8 is translocated into the nucleus in a STAG3-dependent manner. REC8/STAG3-containing cohesin regulates chromosome topological properties and specifically maintains centromeric cohesion. Second, REC8 and mitotic cohesin RAD21 are located at adjacent sites but predominantly at nonoverlapping sites on ESC chromosomes, implying that REC8 can function independent of RAD21 in ESCs. Third, knockdown of REC8-cohesin not only leads to higher rates of premature centromere separation and stalled replication forks, which can cause proliferation and developmental defects, but also enhances compaction of the chromosome structure by hyperloading of retinoblastoma protein condensin complexes from prophase onward. We propose that the delicate balance between mitotic and meiotic cohesins may regulate ESC- specific chromosomal organization and mitotic program.

## Introduction

Embryonic stem cells (ESCs) undergo global chromosome decondensation and dynamic changes in the chromatin landscape to express diverse genes upon lineage differentiation (Liu *et al*, 2019; Dixon *et al*, 2015). ESCs and differentiated cells are notably different; for example, ESCs have self-renewal capabilities, express high levels of DNA repair proteins, and proliferate rapidly with a prolonged S phase during the cell cycle (Zwaka & Thomson, 2003; Choi *et al*, 2017; Choi *et al*, 2018; Choi *et al*, 2020). As such, ESCs display a high tolerance for endogenous replication and DNA damage stress (Choi *et al*, 2017; Ahuja *et al*, 2016; Vitale *et al*, 2017). However, little is known about how ESCs maintain genome integrity and cope with the chromosomal abnormalities and replication stresses that may occur during cell proliferation and early embryogenesis.

Cohesion between sister chromatids allows precise chromosome morphogenesis, recombination, and transcriptional regulation (Kagey *et al*., 2010; Nasmyth & Hearing 2009; Peters *et al*, 2008; Onn *et al*, 2008). Although cohesin was initially identified for its role in sister chromatid cohesion, cohesin is involved in a variety of cellular processes, including DNA replication, chromosome segregation, and DNA damage repair (Peters *et al*, 2008; Wu & Yu 2012; Litwin *et al*, 2018; Hirano 2016; Michaelis & Nasmyth 1997; Watrin & Peters 2009; Schöckel *et al*, 2011). Cohesin is therefore an important candidate for managing topological changes of chromosomes and maintaining genome integrity in ESCs. Cohesin forms V-shaped structures that wrap the sister chromatids together from S phase to anaphase (Gruber *et al*, 2003; Zhang *et al*, 2008) (Fig 1A). Most eukaryotic cells express two α-kleisin proteins, RAD21 and REC8. RAD21 is expressed in both mitotic and meiotic cells. In contrast, REC8, a RAD21 paralog, is expressed in only meiotic cells (Peters *et al*, 2008). In mammals, RAD21 binds to the two orthologous STAG subunits, STAG1 and STAG2, which further interact with a heterodimer composed of WAPL and PDS5 (Gandhi *et al*, 2006; Zhang *et al*, 2006). The cohesin complex consists of 50-nm coiled-coil SMC subunits (SMC1A and SMC3 for mitosis; SMC1B and SMC3 for meiosis) that join together to form a hinge domain and an α-kleisin subunit that interacts with diverse cohesin subunits (Nasmyth & Hearing 2009). In the meiotic cohesin complex, SMC1A and RAD21 are replaced by SMC1B and REC8, respectively (Nasmyth & Hearing 2009). Other cohesin subunits also have distinct mitotic and meiotic isoforms. For example, STAG1 and STAG2, cohesin accessory subunits, can be replaced with the meiotic factor STAG3.

**Figure 1.**
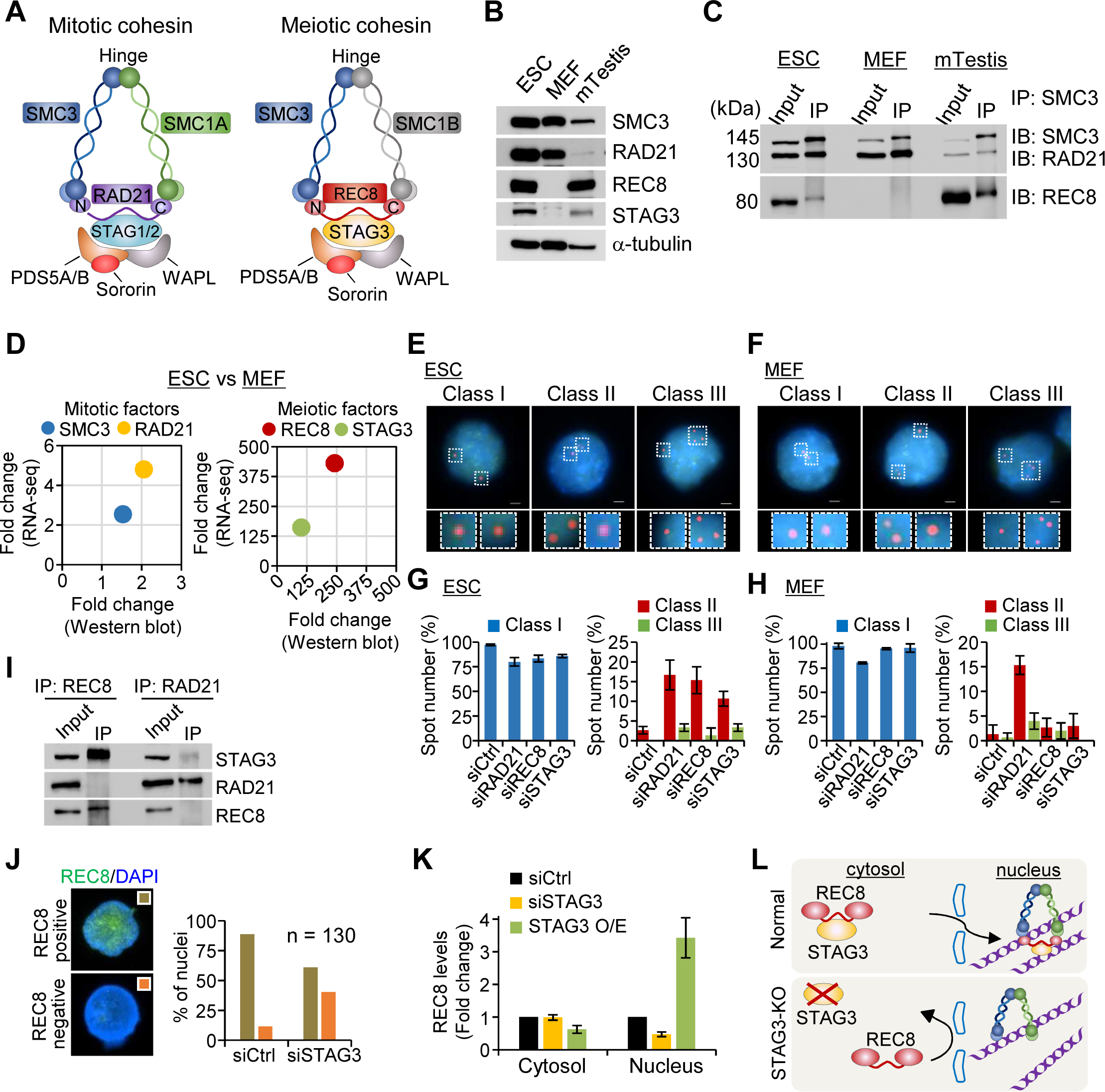
Expression and localization of meiotic cohesin components in ESCs. (**A**) Cohesin complexes in mitosis and meiosis. Cohesin complex is composed of long coiled-coil SMC subunits, called SMC3 and SMC1A for mitotic cohesins, and SMC3 and SMC1B for meiotic cohesins (Gruber *et al*, 2003; Zhang *et al*, 2008). SMC complexes combined with the α-kleisin subunits (RAD21 for mitotic cohesion and REC8 for meiotic cohesin) form a V-shaped structure (Gruber *et al*, 2003; Zhang *et al*, 2008). Cohesin subunits also include STAG1/2 for mitotic cohesin, STAG3 for meiotic cohesin, PDS5A/B, Sororin, and WAPL, which are directly involved in the formation of cohesin complexes. (**B**) Expression analysis of cohesin components. Whole-cell lysate samples were extracted from asynchronous ESCs (J1), MEFs, and testis of C57BL/6J mice. α- tubulin was used as a loading control. (**C**) Immunoprecipitation (IP) analysis in ESCs, MEFs, and mouse testis. We pulled down the cohesin subunits using anti-cohesin (SMC3, RAD21, and REC8) antibodies from whole-cell extracts in ESCs, MEFs, and mouse testis. (**D**) Comparison of expression level of SMC3, RAD21, REC8, and STAG3 between ESCs and MEF, as assessed by Western blotting and RNA-sequencing analyses. (**E** and **F**) Representative images of the interphase FISH analysis for the detection of sister-chromatid separation in ESCs and MEFs. The signals of the locus-specific probe, bound to arm regions (Chromosome 4: 116,094,264 – 116,123,690), are shown in red and DAPI-counterstained chromosomes are shown in blue. Scale bars are 2.5 μm. (**G** and **H**) Quantification of spots in each nucleus. At least 150 nuclei were counted for each experiment. Error bars represent mean ± SD value from three independent experiments. (**I**) Physical interaction between REC8 and STAG3 in ESCs as demonstrated by IP analysis. (**J**) Quantification of REC8-positive nuclei in siCtrl and siSTAG3 (n = 130). (**K**) Quantification of REC8 localization based on the expression pattern of STAG3 in the cytosol and nucleus. Error bars are the mean ± SD. (**L**) Translocation of the cytosolic REC8–STAG3 complexes into the nucleus by direct physical interaction. Under the normal condition, STAG3 directly binds to REC8 and translocates REC8 into the nucleus. However, after STAG3 knockdown, REC8 localization into the nucleus is defective.

Condensin has also been implicated in chromosome organization and morphogenesis (Hirano, 2016). Condensins are ring-shaped protein complexes that mediate chromosome compaction and segregation during mitosis and meiosis (Hirano 2016; Yu & Koshland 2005; Coelho *et al*, 2003). Recent studies have demonstrated that condensins are involved in a wide range of chromosome-related aspects and processes such as genome integrity, genetic recombination, epigenetic regulation, and differentiation (Hagstrom & Meyer 2003; Hirota *et al*, 2004). It has been previously reported that retinoblastoma-associated protein (RB) is required to facilitate chromosome condensation by recruiting condinsin complex to chromosomes (Coschi *et al*, 2010; Coschi *et al*, 2014). Thus, RB is generally thought of as a condensin modulator in the chromosome condensation progression. Most eukaryotes have two types of condensin complexes, condensin I and condensin II; their core subunits SMC2 and SMC4 belong to the chromosomal ATPase family called the structural maintenance of chromosomes (SMC) family, which is conserved in most eukaryotic species (Hirota *et al*, 2004; Skibbens 2019). Further, the complexes contain a unique set of non-SMC regulatory subunits, HEAT-repeat and kleisin subunits (Hirano 2016; Hirota *et al*, 2004; Hirano *et al*, 1997).

During meiosis, meiotic cohesin components, including REC8 and its interacting subunits, play essential roles in not only sister chromatid cohesion but also genetic recombination and chromosome morphogenesis (Brar *et al*, 2009; Hong *et al*, 2019; Yoon *et al*, 2016). The meiotically expressed SCC1/RAD21 can partially substitute for REC8 to support chromosome axes and interhomolog partner interactions during recombination (Biswas *et al*, 2016; Kim *et al*, 2010). However, REC8 has meiosis-specific activities carried out through its interactions with meiosis-specific proteins (Biswas *et al*, 2016; Kim *et al*, 2010). Furthermore, during meiotic prophase, cohesin associates with the chromosome axis and promotes synaptonemal complex formation, homolog template choice, and crossover-specific recombination (Yu & Koshland 2005; Hong *et al*, 2019; Kim *et al*, 2010; Lee *et al*, 2017). The partial loss of cohesin causes centromere decompaction and kinetochore fragmentation, especially in aged oocytes (Zielinska *et al*, 2019; Gruhn *et al*, 2019). These features imply that both mitotic and meiotic cohesins regulate chromosomal morphogenesis during prophase and support the accurate segregation of anaphase chromosomes. However, it is unknown whether the meiosis-specific cohesin components contribute to mitotic chromosome organization.

The present study addresses unexpected phenomena related to the dynamic activity of meiosis-specific cohesin components, REC8 and STAG3, in mitotic ESC chromosomes. We first observed that ESCs express high levels of both REC8 and STAG3. Thus, we sought to determine whether these meiotic cohesion components play distinct and overlapping roles in the presence of mitotic cohesins in chromosome structure organization, sister cohesion, and DNA replication in ESCs. In this study, our data demonstrated multiple functions of the meiosis-specific cohesin components REC8 and STAG3, along with mitotic cohesin subunit RAD21, expressed in ESCs. STAG3 induces the translocation of REC8 into the nucleus, indicating that the REC8/STAG3- containing cohesin complex is functionally active in ESCs. We also show that REC8 could compete with RAD21 for the structural organization of tightly conjoined sister chromosomes and specifically maintains centromeric cohesion, in addition to its general role in cohesion. High-resolution 3D imaging revealed that after knockdown of mitotic or meiotic cohesins, hyperloading of RB-condensin complexes on chromosomes induces the formation of short and fat chromosomes. Mechanisms of collaboration and competition between mitotic and meiotic cohesins provide a new view of how two different cohesin complexes regulate the topology of mitotic chromosomes and sister chromatid cohesion in ESCs.

## Results

### Expression of REC8 and STAG3 in ESCs

Unlike differentiated cells, ESCs exhibit a globally open chromatin structure with various epigenetic landscapes (Dixon *et al*, 2015; Choi *et al*, 2017; Maia *et al*, 2011), which might explain why ESCs can express diverse ESC-specific genes involved in stemness and proliferation (Tee & Reinberg 2014). We therefore aimed to determine whether meiotic cohesin components could be expressed or function as mitotic cohesin components in mouse ESCs. We first analyzed the expression pattern of cohesin proteins in whole-cell lysates isolated from ESCs, mouse embryonic fibroblasts (MEFs), and testes from mice by western blotting. The mitotic cohesin components SMC3 and RAD21 were expressed in both ESCs and MEFs; remarkably, the meiosis-specific cohesin components REC8 and STAG3 were abundantly expressed in ESCs but not in MEFs (Fig 1B; Supplementary Figure S1). Since ESCs expressed high levels of REC8 and STAG3, we concluded that these proteins might play a role in sister chromatid cohesion in ESCs.

To further determine whether the meiotic cohesin components are linked to the SMC ring and α-kleisin subunit, which connects the long coiled-coil arms of the ring, or forms an intact cohesin complex with mitotic components, we performed co-immunoprecipitation (co-IP) assays. We analyzed whether SMC3 forms a complex with RAD21, REC8, and STAG3 by using antibodies against RAD21, REC8, and STAG3. As shown for RAD21, REC8 and STAG3 also strongly interacted with SMC3 in ESCs and testis tissues (Fig 1C; Supplementary Figure S2A). This indicates that REC8/STAG3-containing cohesin may have various functions in the mitotic program of ESCs.

### REC8 and STAG3 are required to ensure faithful chromosome segregation in ESCs

Cohesins play a role in sister chromatid cohesion and thus ensure faithful chromosome segregation during mitosis (Gruber *et al*, 2003; Zhang *et al*, 2008; Haarhuis *et al*, 2014; Weitzer & Uhlmann 2002). Therefore, we aimed to determine whether the meiotic cohesin components REC8 and STAG3 mediate proper cohesion, similar to mitotic cohesins, in ESCs and MEFs. We depleted the mitotic cohesin component RAD21, and the meiotic cohesin components REC8 and STAG3 by treatment of cells with pooled siRNAs against STAG3, RAD21, and REC8 (siSTAG3, siRAD21, and siREC8). In ESCs and MEFs, the knockdown efficiency of the cohesin gene-targeted siRNA pool was more than 60% greater than that of siControl (siCtrl) (Supplementary Figure S2B), and the decreased expression of individual cohesin component genes did not affect the expression levels of the other cohesin components (Supplementary Figure S2B). Expression levels of RAD21, REC8, and STAG3 were higher in ESCs than in MEFs: RNA-seq analysis confirmed that changes in cohesin expression levels were similar to those observed by protein blot (Fig 1D).

Interphase analysis using fluorescence in situ hybridization (FISH) in cells with siRNA-mediated knock down of mitotic or meiotic cohesin components was used to measure the level of sister chromatid separation, which can be identified by the number of distinct foci corresponding to a given locus; we hybridized a locus-specific probe (chromosome 4) to interphase ESCs, performed BrdU incorporation, and counted the number of fluorescence signals from the hybridization sites of two homologous sequences. Under siCtrl treatment, interphase cells showed Class I signals (two spots), which corresponded to the two sister chromatids. Upon cohesin component knockdown, the incidence of abnormal sister chromatid separation (Class II (three spots) and III (four spots)) was significantly increased, ranging from 14–20% in ESCs lacking RAD21, REC8, or STAG3 (Fig 1E and G). In MEFs, RAD21 knockdown showed a higher rate of abnormal sister chromatid separation (∼ 20%) than REC8 (∼ 4%) and STAG3 (∼ 3%) knockdown (Fig 1F and H). Overall, REC8 and STAG3 knockdown resulted in significant sister chromatid cohesion defects. These results support our hypothesis that REC8 and STAG3 are involved in mitotic chromosome cohesion to ensure faithful chromosome separation in ESCs.

### STAG3 induces the translocation of REC8 into the nucleus by interacting with REC8

STAG3 mediates REC8 stabilization, and the REC8–STAG3 complex is required for diverse cohesin functions, chromosome axis organization, and recombination during meiosis (Fukuda *et al*, 2014; Wolf *et al*, 2018). To investigate whether STAG3 is required for the role of REC8 in ESCs, we characterized the REC8-STAG3 interaction using a protein immunoprecipitation assay. STAG3 physically interacted with REC8 but not with RAD21 (Fig 1I), implying that REC8 and STAG3 can cooperate to promote the formation of cohesin complexes. To better understand the localization of REC8 and STAG3, we analyzed REC8 positive and negative signals in nuclei by immunofluorescence microscope. REC8-negative nuclei were significantly increased by siSTAG3 (40.2%) compared to those in siCtrl nuclei (11.3%). (Fig 1J). Further, fractionation of cytosol and nuclear proteins from ESCs revealed that REC8 and STAG3 were present in both the nucleus and the cytosol. Upon STAG3 knockdown, the REC8 level in the nucleus was reduced by ∼52%. However, overexpression of STAG3 showed greater levels of REC8 in the nucleus more than 3.4- fold compared to siCtrl (Fig 1K; Supplementary Figure S2C, D). Thus, we suggest that REC8 is translocated into the nucleus in a STAG3-dependent manner (Fig 1L) but did not affect the translocation of SMC3 or RAD21.

### The role of REC8–STAG3 cohesin complexes in replication fork progression in ESCs

Sister chromatid cohesion is established during DNA replication in both mitosis and meiosis (Peters, 2012; Sherwood, 2010). As cohesin components are enriched at the DNA replication fork and provide a bridge linking the sister chromatids (Peters *et al*, 2008; Sherwood *et al*, 2010; Guillou *et al*, 2010), we explored whether the REC8–STAG3 cohesin complexes play a role in the DNA replication process. To analyze the molecular functions of REC8–STAG3 cohesin complexes in DNA replication, we consecutively pulse labeled ESCs and MEFs with the nucleotide analogs CldU and ldU and observed the DNA replication process in a single strand of DNA (Fig 2A and D). The labeled elongating DNA strands were immunostained with antibodies to visualize the progression of the replication forks. The track length and fork progression rate were significantly decreased by more than 1.4-fold in SMC3-, RAD21-, REC8-, and STAG3-knockdown ESCs compared with siCtrl ESCs (Fig 2B and C), which implied an increase in the frequency of stalled replication forks. In MEFs, the length of replicating DNA strands and the rate of DNA replication elongation after knockdown of cohesin components decreased by 1.45-fold and 1.48-fold with siSMC3 and siRAD21, respectively; however, we did not observe any significant difference after the knockdown of REC8 (1.04-fold) or STAG3 (1.01-fold) (Fig 2E and F). We further analyzed the cell cycle and found that downregulation of cohesin led to slowed S-phase progression in ESCs (Supplementary Figure S3).

**Figure 2.**
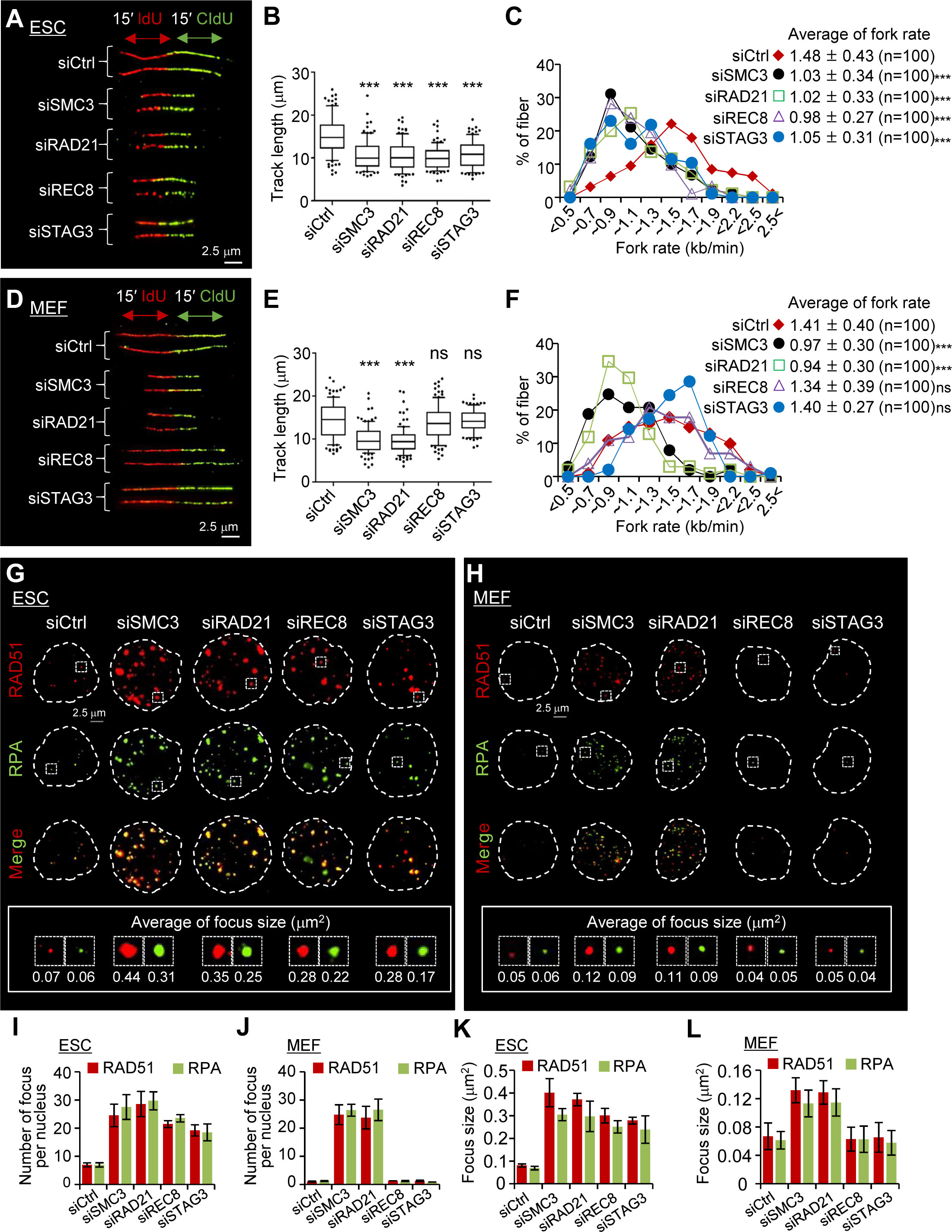
The role of meiotic cohesins in replication fork progression. (**A** and **B**) DNA fiber labeling scheme (Top): IdU is incorporated at the first analog (15 min), followed by CldU (15 min), incorporated as the second analog. Representative image of DNA fiber in ESCs and MEFs (bottom). Scale bars are 2.5 μm. (**B** and **E**) Length of IdU and CIdU tracks in siCtrl and siCohesin. We transfected the cells with siCtrl or siCohesin (siSMC3, siRAD21, siREC8, and siSTAG3) before DNA labeling, as indicated. Top and bottom dots of the box/whisker plot represent high and low rank 10% and the box plot indicate lower quartile in which seventy-five percent of the scores fall below the upper quartile value, median, which average value of all data, and low quartile, which twenty-five percent of scores fall below the lower quartile value. In each experiment, we analyzed at least 50 independent fibers. Results are illustrated as ± SD value. Statistics: Mann– Whitney t-test, ns: not-significant, ***P < 0.001. (**C** and **F**) Elongation rates during DNA replication. We measured the lengths of the DNA fibers (IdU and CldU tracks) as shown in (A). To calculate the DNA fork rates, we converted the DNA fiber lengths to kilobase (kb) (1 mm = 2.94 kb) and converted values divided by CldU/IdU pulse labeling time (30 min). P-values were calculated by Student’s t-test and n is the number of measured fibers. (**G** and **H**) Images of RAD51 and RPA foci formation in ESCs and MEFs (top). Focus size in each condition was analyzed using Nikon software (bottom). Scale bars are 2.5 μm. (**I** and **J**) Numbers of RAD51 and RPA foci were counted in siCtrl (control) cells and siCohesin (siSMC3, siRAD21, siREC8, and siSTAG3)-treated cells. At least 50 nuclei were counted for each condition. Error bars represent mean ± SD value from three independent experiments. (**K** and **L**) Average of focus size after knockdown of cohesin components. Focus size was counted in normal cells and cohesin knockdown cells. At least 50 nuclei were counted for each experiment. Error bars represent mean ± SD value from three independent experiments.

These results suggest that the REC8–STAG3 cohesin complexes are involved in the progression of DNA replication in ESCs.

Based on these data, we further hypothesized that the REC8–STAG3 cohesin complexes might also participate in the DNA gap repair process to maintain genome integrity. We analyzed the formation of RAD51 and replication protein A (RPA) foci after knocking down the targeted cohesin components in ESCs and MEFs with siSMC3, siRAD21, siREC8, and siSTAG3. Knockdown of SMC3, RAD21, REC8, and STAG3 dramatically increased the number of RAD51 and RPA foci in ESCs but not in MEFs (Fig 2G and H). REC8 and STAG3 knockdown did not affect the number of RAD51 or RPA foci (Fig 2I and J). Moreover, the size of the foci in ESCs was also considerably increased compared to that in MEFs (Fig 2G and H; Fig 2K and L). Thus, the lack of cohesion established by the REC8–STAG3 cohesin complexes could stall DNA replication and RAD51 and RPA foci accumulation at DNA breaks. We further found that knockdown of REC8 resulted in a significant increase in the proportion of apoptotic cells among ESCs (Supplementary Figure S4), suggesting that the REC8–STAG3 cohesin complexes contribute to cohesion-mediated cellular progression.

### Knockdown of REC8 and STAG3 causes precocious sister chromatid separation and hypercompaction of chromosomes in ESCs

Cohesin plays an important role in the protection of centromeric cohesion during both mitosis and meiosis (Peters *et al*, 2008; Diaz-Martinez *et al*, 2007; Carretero *et al*, 2013). To investigate whether the REC8–STAG3 cohesin complexes play a role in the maintenance of sister chromatid cohesion in ESCs, we used FISH assays with a telomere DNA probe to analyze chromosome cohesion in metaphase chromosome spreads. Knockdown of cohesin components in ESCs resulted in increased rates of precocious sister chromatid separation (28.03% with siRAD21, 24.64% with siREC8, 15.98% with siSTAG3, and 4.20% with siCtrl) (Fig 3A–D). In MEFs, the rate of precocious centromere separation was increased by siRAD21 (27.28%) but not by siREC8 (6.15%) or siSTAG3 (7.12%) compared to that in siCtrl cells (4.02%) (Fig 3A–D).

**Figure 3.**
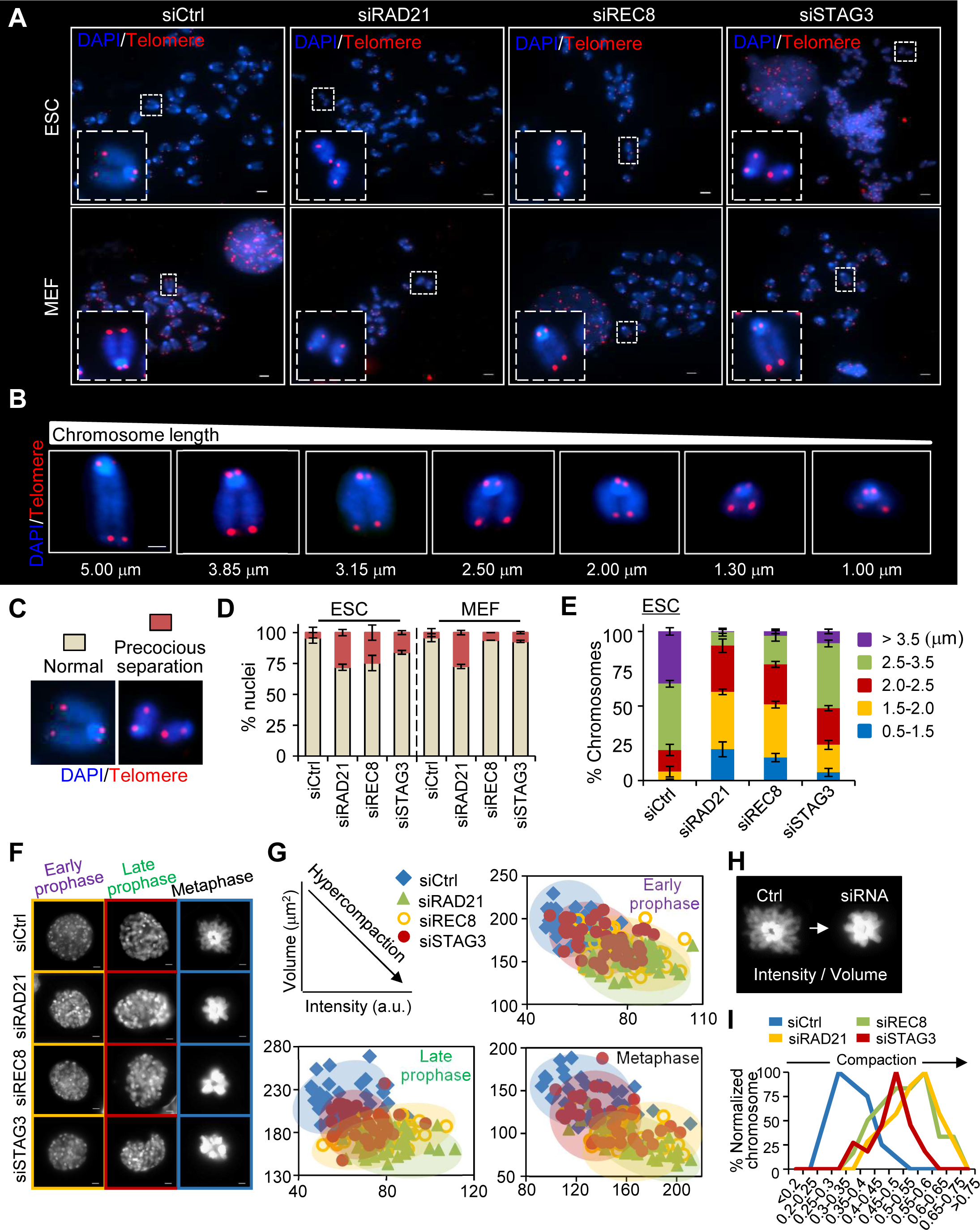
Chromosome compaction and precocious separation after the knockdown of cohesin components. (**A**) Metaphase chromosome images of telomere FISH from ESCs and MEFs expressing a non-targeting siRNA or different siRNAs against RAD21, REC8, and STAG3 (siRAD21, siREC8, and siSTAG3). Telomeric probe signals are shown in red and DAPI-counterstained chromosomes are shown in blue. Scale bars are 10 μm. (**B**) Metaphase chromosome images of diverse length. Scale bar is 2.5 μm. (**C**) Images of a metaphase spread from normal (left) and precocious separation (right) showing proper and defective chromosome separation, respectively. (**D**) Quantification of normal cells and precocious separated cells. The bar graph indicates the percentages of cells showing a ratio of normal chromosomes and precocious separated chromosomes among n ≥ 200 metaphase cells in each condition. Error bars represent mean ± SD value from three independent experiments. (**E**) Quantification of chromosome length and percentage of average chromosome length per metaphase cells. Length of chromosomes was measured by calculating the distance between both sides of the telomere probes. Data are shown as mean ± SD value from at least 200 chromosomes per condition. (**F**) Compaction analysis of chromosomes from prophase to metaphase. ESCs expressing histone H2B-GPF were analyzed by fluorescence microscopy. Scale bars are 2.5 μm. (**G**) Level of chromosome compaction from prophase to metaphase. We analyzed cell volume and intensity using the Nikon NIS software. X and Y axes of the graphs indicate intensity and volume (μm2), respectively. (**H**) Representative images of ESCs marked with H2B-GFP in metaphase. (**I**) Quantification of the compact level of chromosomes. The level of compaction is calculated by dividing the GFP-signal intensity with the chromosome volume. At least 30 metaphase cells were counted per condition.

To evaluate changes in length distributions, we classified the chromosome length into five numerical ranges and also described the length by tracking the distance between the telomeres of sister chromatids. In ESCs lacking RAD21, REC8, and STAG3, the proportion of chromosomes with lengths in the 0.5–2.5 µm range were approximately 70%, 57.5%, and 28% higher than those of siCtrl cells, respectively (Fig 3B and E); on the other hand, in siCtrl cells, the proportion of chromosomes with lengths over 2.5 µm was 70%, 57.5%, and 28% higher than that in siRAD21, siREC8, and siSTAG3 cells, respectively. In contrast, the chromosome length distributions in MEFs did not change significantly except after the knockdown of RAD21 (Supplementary Figure S5A). Thus, when meiotic/mitotic cohesin components were depleted, chromosome hypercompaction occurred in ESCs (Supplementary Figure S6B), which also suggests that the REC8–STAG3 cohesin complexes have overlapping roles with RAD21-containing mitotic cohesin complexes in the organization of chromosome topology during prophase and metaphase in ESCs.

Evidence suggests that cohesin contributes to chromosome compaction during mitosis (Peters *et al*, 2008; Hirano 2016; Guillou *et al*, 2010). However, it remains unclear how chromosome compaction is regulated in normal or aberrant sister chromatid cohesion. To understand the role of cohesin in chromosome morphogenesis in ESCs, we next examined its effects on chromosome volume and intensity from early prophase to metaphase by stably expressing GFP-labeled histone H2B (H2B-GFP) in these cells. The chromosomes were more dramatically compacted in cells treated with siRNA targeting cohesin components than in siCtrl cells, becoming both denser and fatter from early prophase to metaphase (Fig 3A, F, and G). Additionally, the intensity/volume values were increased after siRAD21, siREC8, and siSTAG3 treatment compared to siCtrl treatment (Fig 3H and I).

To measure chromosome compaction more precisely, we performed FISH analysis using two probes to mark telomeres and a locus in the arm of chromosome 4 (chromosome 4: 116,094,264–116,123,690) and determined the distance between each telomere and the specific locus in the arm. In ESCs lacking RAD21, REC8, and STAG3, the long and short distances between each telomere and the locus in the arm were reduced by more than 1.1 µm and 0.5 µm, respectively (Fig 4A–C). In contrast, the chromosome distance in MEFs was not changed significantly except after RAD21 knockdown (Fig 4A–C). Taken together, we conclude that the loss of cohesin components induced irregular chromosome compaction at metaphase.

**Figure 4.**
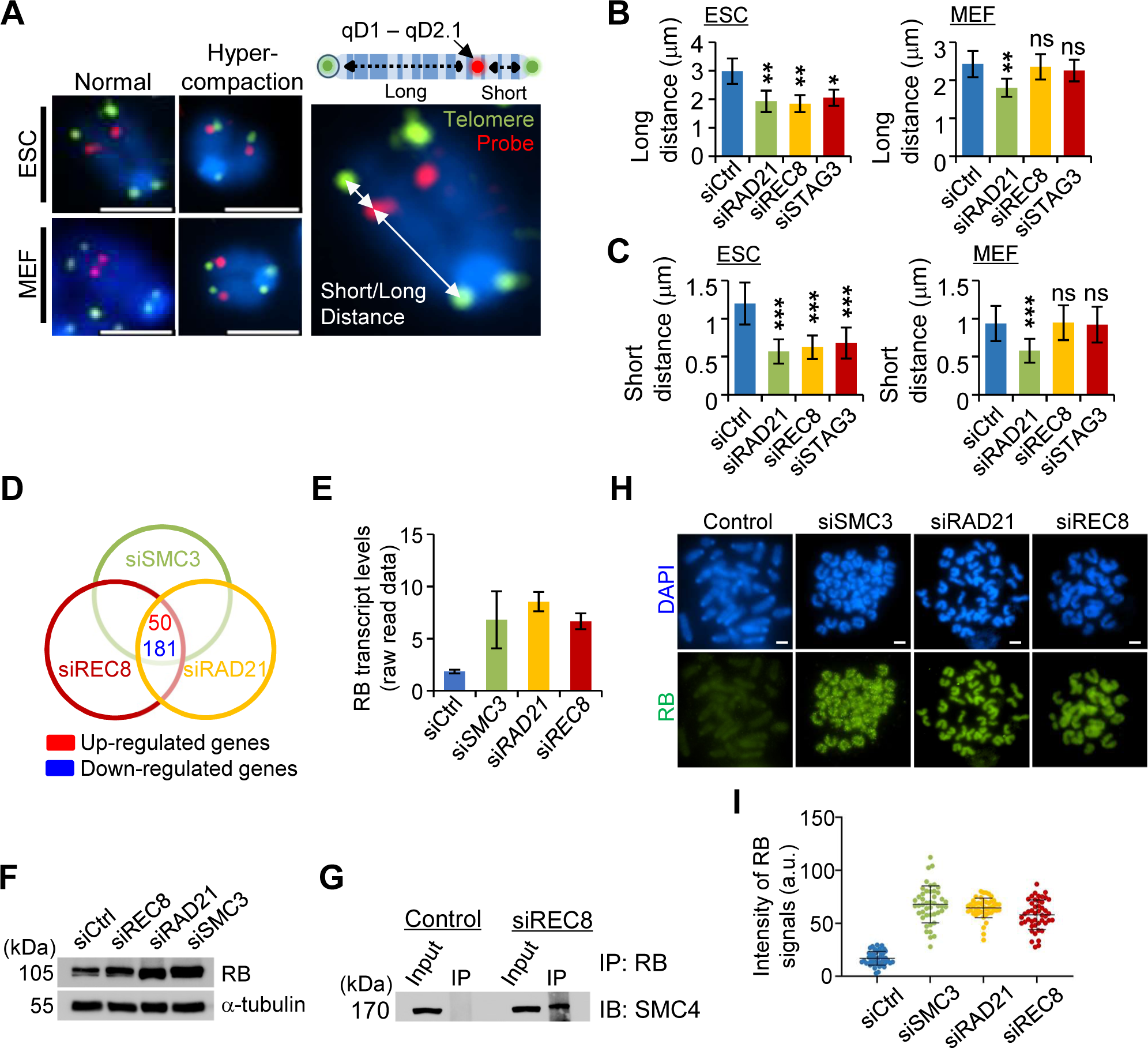
Chromosome shortening at metaphase chromosome in the knockdown of cohesin components. (**A**) Metaphase spreads hybridized with the telomeric probe and the locus-specific probe that is bound to the region of Ch 4 (Chromosome 4: 116,094,264–116,123,690). Telomeric probe and locus-specific probe signals are shown in green and red, respectively. The lengths of chromosomes were measured by calculating the distance between the telomere and locus-specific probes. The chromosome lengths were analyzed using Nikon NIS software. Scale bars are 2.5 μm. (**B and C**) Quantification of chromosome lengths in metaphase chromosomes from cells treated or untreated with siRNA specific to the cohesin components for 24 h. We measured the short or long chromosome distance in ESCs and MEFs. Error bars represent mean ± SD value from three independent experiments (at least 30 metaphases per condition). ns: not-significant, *P < 0.05 and **P < 0.01 (Student’s t-test) are significant compared to siCtrl (Control). (**D**) A Venn diagram for identification of up- or down-regulated genes from siRNA-mediated knockdown of SMC3, RAD21, and REC8 in ESCs. Data were adjusted with P < 0.1 and fold change > 1.5. (**E**) A comparison of gene expression of RB. RB transcript levels from raw read data of RNA-sequencing experiments were analyzed to evaluate RB gene expression (Supplementary table 1). (**F**) Analysis of RB protein levels following transfection with a siRNA pool against REC8, RAD21, and SMC3 in ESCs. α-tubulin used as a loading control. (**G**) Immunoprecipitation analysis for physical interaction between RB and SMC4 in ESCs. (**H**) Immunofluorescence analysis of chromosome compaction in metaphase of ESCs. Chromosomes were stained with anti-RB antibody and DAPI. Control, siControl; siRAD21 or siREC8, siRNA treatment against RAD21 and REC8, respectively. Scale bars are 2.5 μm. (**I**) Intensity of RB protein signals per a chromosome in ESCs. a.u., arbitrary unit. Results represent as mean ± SD value (N > 50).

### Elevated expression of RB promotes chromosome hypercompaction

To investigate global changes in gene expression in response to aberrant sister chromatid cohesion that can be used to identify potential targets for chromosome hypercompaction, we performed RNA-Sequencing analysis following treatment with siSMC3, siRAD21, and siREC8. Venn diagram revealed that 50 genes were more highly expressed in cohesin knockdown cells (Fig 4D; Supplementary Table 1). Gene sets of RNA-sequencing and western blotting were used to investigate whether cohesin-knockdown ESCs exhibit changes in expression of RB, a binding partner of condensin. Interestingly, the expression levels of RB were significantly increased in response to SMC3, RAD21, and REC8 knockdown (Fig 4E and F). In addition, through immunoprecipitation of RB followed by western blotting for a condensin component, SMC4, we confirmed that high level expression of RB resulted in a strong increase of RB-condensin interaction after REC8 knockdown (Fig 4G). To determine the consequence of high levels of RB, localization of RB and chromosome compaction were analyzed after transfection with siSMC3, siREC8, and siRAD21. Knockdown of cohesin components in ESCs resulted in increased levels of RB-staining signals in metaphase chromosomes (4-fold in siSMC3, 3.8-fold in siRAD21 and 3.4-fold in siREC8) (Fig 4H and I). Furthermore, the chromosomes were more dramatically compacted in cells transfected with siRNA targeting cohesin components than in siCtrl cells (Fig 4H). These results suggest that aberrant sister chromatid cohesion confers elevated expression of RB and that the abundance of RB-condensin complexes actively promotes hypercompaction of chromosomes.

### REC8 can function independent of RAD21 in mitotic ESC chromosomes

Cohesins present at the interface of sister chromatids are linked by DNA/structural bridges in prophase (Chu *et al*, 2020; Giménez-Abián *et al*, 2004); thus, both REC8 and RAD21 might be independently localized between sister chromatids. We performed microscopic analysis of REC8 and RAD21 to detect cohesin localization on metaphase chromosomes. WAPL promotes the dissociation of cohesin complexes from chromosomes by providing a DNA exit gate during prophase and metaphase (Gandhi *et al*, 2006; Haarhuis *et al*, 2017). We therefore suppressed the expression of WAPL, a cohesin release factor, and/or REC8 and RAD21 using siRNA and then investigated the localization of mitotic RAD21 cohesin and meiotic REC8 cohesin on metaphase chromosomes by immunofluorescence microscope. We found that RAD21 and REC8 were highly accumulated on closed chromosomes after WAPL knockdown (Supplementary Figure S6A). The rate of closed chromosome morphology was higher in cells with WAPL knockdown (74.9%) than in siCtrl cells (10.8%) (Supplementary Figure S6B), indicating that chromosome arm-bound cohesins were stably maintained.

We then analyzed the specificity in binding patterns of RAD21 and REC8 during metaphase by immunofluorescence microscope. After WAPL knockdown, RAD21 and REC8 were located at adjacent sites but predominantly at nonoverlapping sites on chromosome arms (Fig 5A). Thus, REC8 likely operates independently of RAD21 and complements its function. We then measured the cohesin intensity following the co-depletion of WAPL and RAD21 or REC8. When both RAD21 and WAPL were depleted, the intensity of REC8 on chromosomes increased more than 2.03-fold compared with that in cells with knockdown of WAPL alone (Fig 5A and B; right panel). Additionally, we showed that RAD21 intensity was increased more than 1.34-fold in cells depleted of both WAPL and REC8 compared with WAPL-knockdown cells (Fig 5B; left panel), suggesting that RAD21 may compete with REC8 to form cohesin complex. Additionally, we more precisely analyzed the binding regions of RAD21 and REC8 with high-resolution 3D SIM imaging. The two proteins were found to localize at discrete sites along the chromosome axis and the adjacent regions, but they rarely overlapped (Fig 5C). In particular, REC8 but not RAD21 strongly accumulated in the centromere region (Fig 5D and E; Supplementary Figure S6C). To better characterize REC8 localization in the centromere regions, we immunostained cells for centromere protein C (CENP-C), and found that REC8 colocalized with CENP-C (Fig 5F), further confirming that REC8 is localized to the centromere region. Furthermore, ChIP analysis of the centromere satellite sequences revealed that REC8 was more abundant than RAD21 in centromere regions (Fig 5G). Thus, REC8 could specifically function to support centromeric cohesion in ESCs.

**Figure 5.**
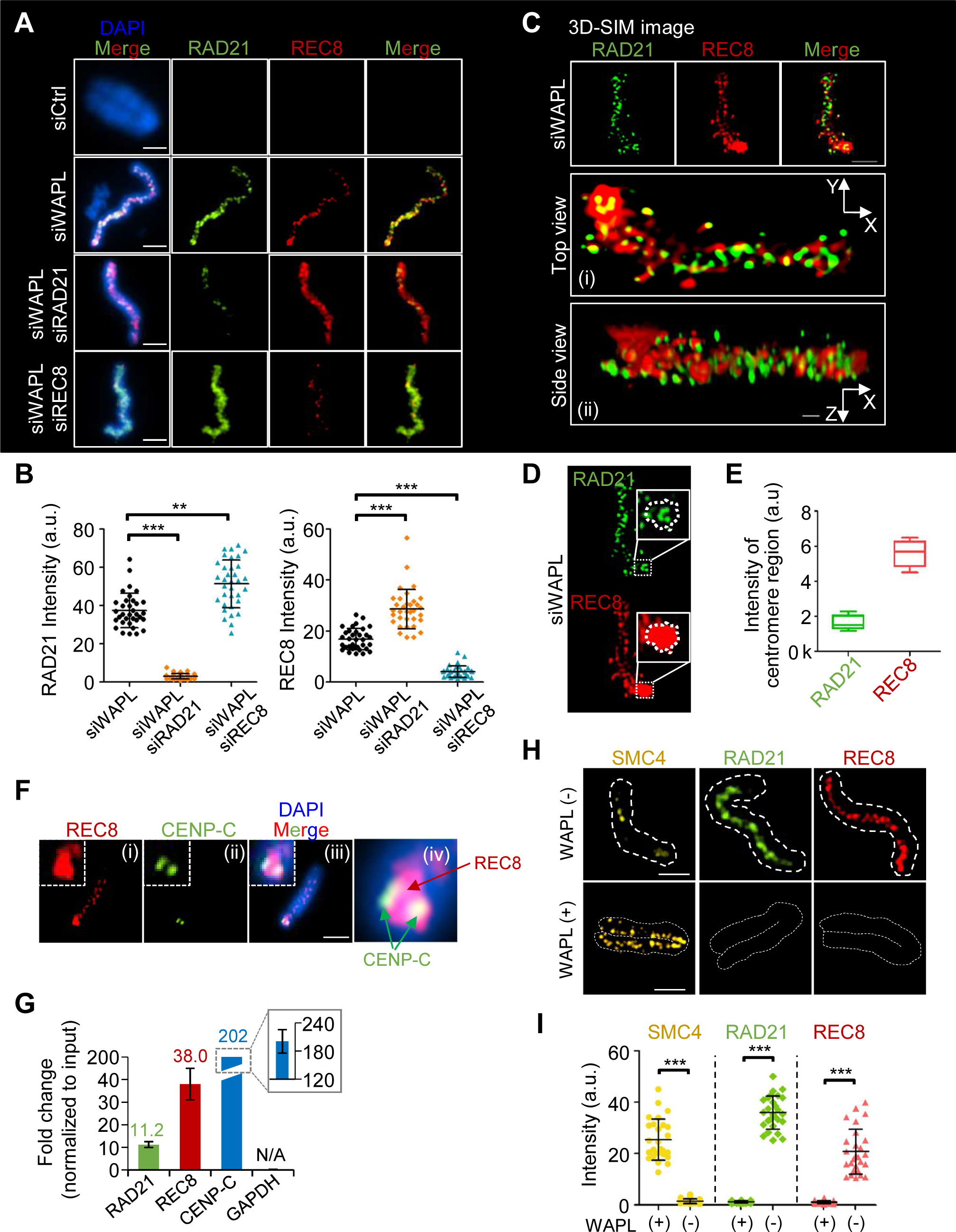
Chromosome association of mitotic and meiotic cohesin components. (**A**) Representative images of WAPL-knockdown cells in metaphase. The cells were treated with siCtrl (Control; first row) or siRNA specific to WAPL (second row), WAPL/RAD21 (third row), and WAPL/REC8 (fourth row). Chromosome spreads were prepared with 0.1 μg colcemid and stained them with antibodies and DAPI. The scale bars are 2.5 μm. (**B**) Quantification of cohesin intensity shown in (A). At least 60 chromosomes were counted. Results are illustrated as mean ± SD value. We calculated P-values (Student’s t-test) using GraphPad Prism 5 software: **P < 0.01, ***P < 0.001. (**C**) 3D structured illumination microscopy (SIM) images of WAPL-knockdown cells in metaphase. The fixed chromosomes were immunostained with specific antibodies. Scale bar is 2.5 μm. 3D SIM images were analyzed using NIS software from Nikon. Scale bar is 10 μm (Side view). (**D**) Representative 3D SIM images of WAPL- knockdown cells in metaphase. (**E**) Quantification of the relative intensity level of RAD21 and REC8 at centromere regions. (**F**) Analysis of centromeric cohesin localization. Fixed chromosomes (C) were immunostained with antibodies (i), CENP-C (ii), merged with DAPI (iii), and enlarged centromere region (iv). Scale bar is 2.5 μm. (**G**) ChIP analysis for RAD21, REC8, CENP-C (positive control of minor satellite), and GAPDH (negative control). ChIP the input signal. Results are illustrated as the mean ± SD value of three independent experiments. (**H**) Fluorescence microscopy images of WAPL- knockdown cells in metaphase. Fixed chromosomes were immunostained with antibodies. Scale bars are 2.5 μm. (**I**) Quantification of cohesin and condensin intensity. Results are illustrated as mean ± SD value (N > 30). P-values (Student’s t-test) were calculated using GraphPad Prism 5 software.

As WAPL knockdown resulted in localization of RAD21 and REC8 on the chromosomes, we next aimed to identify the localization of condensin, which is normally located on sister chromosomes from prophase to anaphase (Hirota *et al*, 2004). As SMC4 is a core component of condensins, we immunostained cells with an anti-SMC4 antibody to observe the localization of SMC4 on chromosomes. Interestingly, after the loss of WAPL, the intensity of SMC4 was reduced by more than 15.1-fold, whereas the intensities of RAD21 and REC8 increased by 31.3-fold and 19.7-fold, respectively (Fig 5H and I), suggesting that cohesin and condensin could be involved in the organization of chromosome structure through their shared DNA binding sites.

### Changes in chromosome structure and SMC4 localization patterns after knockdown of REC8, STAG3, RAD21, and SMC3

Cohesin complexes are released from chromosome arms during prophase (Peters *et al*, 2008; Waizenegger *et al*, 2000) to allow their localization to the mitotic chromosome axis and linkages between different DNA loops, resulting in overall chromosome axis compaction (Hirano 2016; Kschonsak & Hearing 2015; Goloborodko *et al*, 2016; Cockram *et al*, 2020). We therefore expected that knockdown of cohesin would lead to global condensin localization to chromosomes from prophase to metaphase and cause dramatic changes in chromosome compaction. To determine the distribution of condensin after knockdown of REC8, STAG3, RAD21, and SMC3, we arrested cells in metaphase by colcemid treatment and then observed SMC4 by fluorescence microscopy. In ESCs lacking cohesin factors, chromosomes were more compact than those in siCtrl cells and exhibited greater condensin (SMC4) distribution along the whole chromosome, whereas SMC4 in siCtrl cells was located on the mitotic chromosomes (Fig 6A). Additionally, in cells with knockdown of SMC3, RAD21, REC8, or STAG3, SMC4 proteins were distributed more extensively along mitotic chromosomes and showed stronger signals, as shown in 3D projections of chromosomes (Fig 6B and C). However, knockdown of REC8 or STAG3 did not affect the distribution or intensity of SMC4 in MEFs (Fig 6G–I). In cohesin-knockdown cells, we observed numerous SMC4 puncta in compacted chromosomes, indicating that mitotic/meiotic cohesin knockdown specifically induced the recruitment of condensin complexes. Thus, SMC4 could be involved in chromosome compaction during prophase, and timely dissociation of cohesin is required to promote SMC4 loading on chromosomes.

**Figure 6.**
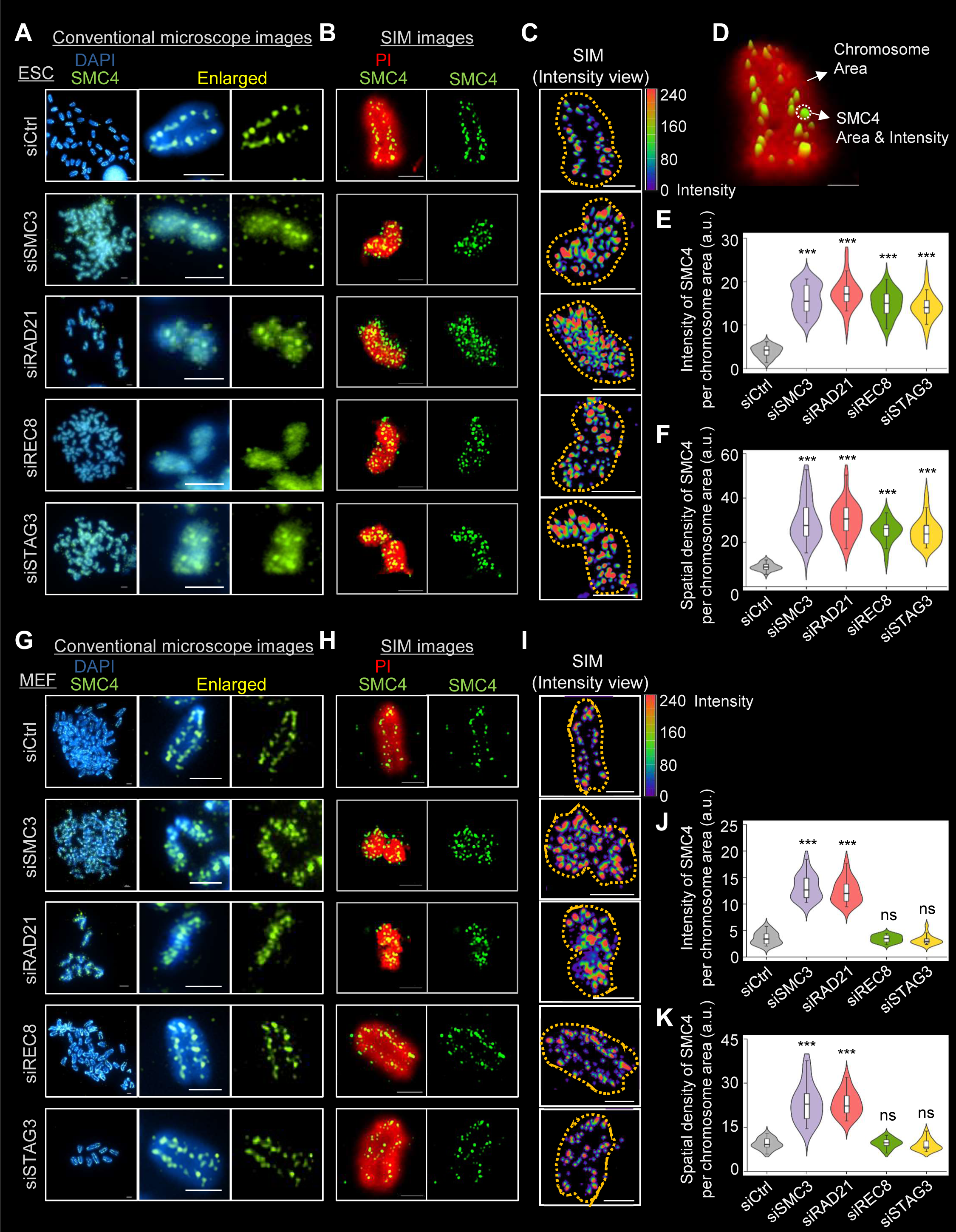
The change in chromosome structures and localization pattern of condensin after knockdown of cohesin components. (**A** and **G**) Fluorescence images of metaphase chromosomes. The chromosomes were immunostained with antibodies against SMC4 and DAPI (left) in ESCs and MEFs. Scale bars are 2.5 μm. (**B** and **H**) 3D SIM images of metaphase chromosomes. We stained the cells with SMC4 and PI. Scale bars are 2.5 μm. (**C** and **I**) 3D-rendering images of SMC4-stained metaphase chromosomes. SMC4 is pseudo-colored in each color to measure the localization and intensity of condensin on metaphase chromosomes. 3D-rendered images were created using the Nikon NIS software. Scale bars are 2.5 μm. (**D**) Intensity and distribution of SMC4 in metaphase chromosomes. (**E** and **F**) Quantification of SMC4 intensity and area in metaphase cells. The area and intensity were measured using NIS software from Nikon. Violin plots show the intensity of SMC4 per chromosome area and the spatial density of SMC4 per chromosome area. The range of the box is from 25%–75% and the black line in each box represents the median. Y-axis values represent the intensity of SMC4 and the spatial density of SMC4 per a chromosome area. Y-axis values were calculated by dividing the volume of chromosome area by the intensity or volume of SMC4 spots. We performed three independent experiments (at least 30 metaphases per condition). P-values were calculated using Student’s t-test. ns: not-significant, *P < 0.5, **P < 0.01, and ***P < 0.001. (**J** and **K**) The intensity and spatial density of SMC4 per chromosome area in MEFs.

To better understand the effects of condensin distribution on cohesin knockdown, we measured the intensity of SMC4 and its spatial density per chromosome area using high-resolution 3D SIM imaging (Fig 6D). The intensity and spatial density of SMC4 after the knockdown of cohesin components were increased by more than 3.7-fold (intensity) and 23% (density) in both ESCs and MEFs. In contrast, compared with siCtrl, siREC8 and siSTAG3 had no significant effect in MEFs (Fig 6E and F; Fig 6J and K).

To determine the cell cycle phase during which chromosomal localization of SMC4 is induced, we next analyzed the intensity of SMC4 during interphase, prophase, and metaphase in ESCs and MEFs. During interphase in ESCs and MEFs, the intensity of SMC4 did not differ between cohesin-knockdown cells and siCtrl cells, implying that SMC4 can initially localize to chromosomes after interphase (Supplementary Figure S7A). In contrast, during prophase and metaphase in ESCs, condensins were distributed in a large area of the chromosomes and more extensively accumulated near the centromeres of cohesin-knockdown cells compared with siCtrl cells (Supplementary Figure S7B, C; top panels). Knockdown of REC8 and STAG3 did not affect the intensity of SMC4 in MEFs during prophase and metaphase (Supplementary Figure S7B, C; bottom panels). Collectively, our data indicate that knockdown of meiotic cohesin induces an irregular distribution of condensins on chromosomes in ESCs, leading to chromosome hypercompaction, and that meiotic cohesins also contribute to the morphogenesis of mitotic chromosomes in ESCs (Supplementary Figure S8A, B).

### Chromosomal loading of SMC4 after knockdown of REC8 and STAG3

As a previous study showed that the binding sites of cohesin complexes overlapped with those of condensin complexes and that they cooccupied the core promoter regions of the ESC pluripotency gene Pou5f1 (Fig 7A) (Dowen, 2013), we speculated that cohesins might compete with condensins at their binding regions and that the knockdown of cohesin could cause condensin complex recruitment to cohesin binding sites. To gain insight into the proportion of the genome occupied by cohesin and condensin complexes in ESCs and MEFs, we performed ChIP analysis with REC8, RAD21, STAG3, SMC3, and SMC4. Interestingly, the level of SMC4 bound at these sites was increased in cohesin component-deficient ESCs compared to siCtrl ESCs by more than 2.24-fold (siSMC3), 2.21-fold (siRAD21), 1.60-fold (siREC8), and 1.58-fold (siSTAG3) (Fig 7B–D). In MEFs, the level of bound SMC4 under cohesin-knockdown conditions was increased by more than 2.89- fold (siSMC3) and 2.41-fold (siRAD21), but knockdown of meiotic cohesin (siREC8: 0.94-fold; siSTAG3: 1.02-fold) did not affect the proportion of bound SMC4 (Fig 7B–D). On the other hand, the levels of SMC3, REC8, and RAD21 bound at these binding sites did not change significantly in ESCs and MEFs (Fig 7B–D). Thus, these results indicate that SMC4 proteins are abundantly loaded at cohesin binding sites after knockdown of cohesin components. These data also imply that REC8– STAG3 cohesin complexes are sufficient to specifically regulate chromosome compaction in ESCs and that the release of these components might enable condensins to bind to cohesin binding sites in ESC chromosomes.

**Figure 7.**
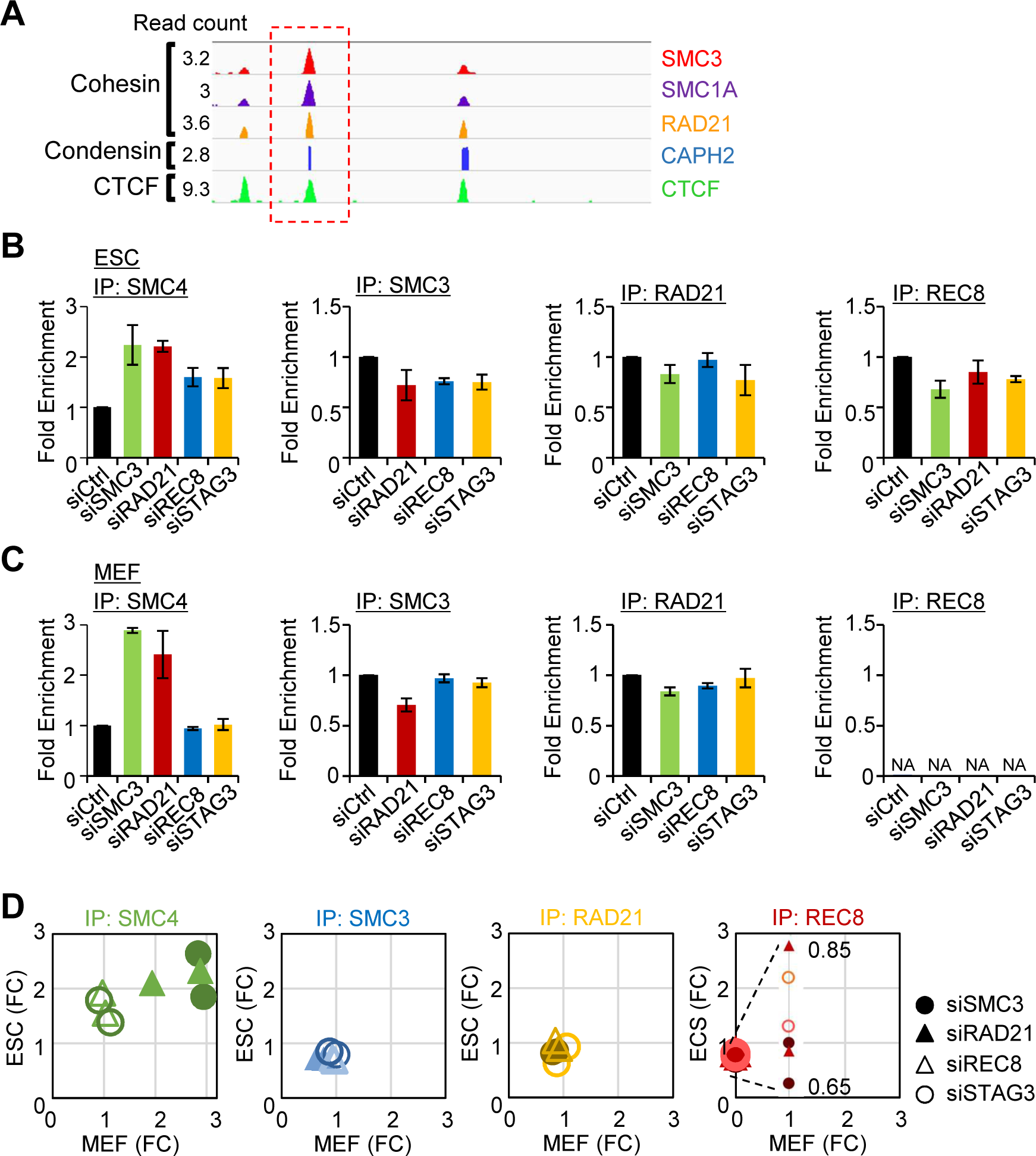
Genome-wide occupancy of cohesin and condensin in ESCs and MEF. (**A**) Binding profiles for cohesin, condensin, and CTCF in ESCs (Dowen *et al*, 2013; Kagey *et al*, 2010). (**B** and **C**) ChIP analysis of mitotic/meiotic cohesin components. We performed ChIP analysis for condensin and cohesin components in ESCs and MEFs. Our data describe the relative level of DNA the pulling down each cohesin component (SMC4, SMC3, RAD21, and REC8) at cohesion and condensin binding regions (Dowen *et al*, 2013). Fold-change values were normalized for each condition with the value of siCtrl (Control) samples. Normalized fold change values in ESCs and MEFs are represented on the bar graphs. Error bars are mean ± SD value. (**D**) Condensin and cohesin ChIP analyses using siRNA in ESCs and MEFs. The data show the relative level of DNA bound in siCtrl (Control) and other siCohesin cells after we pulled-down each cohesin component (SMC4, SMC3, RAD21, and REC8) at the cohesin/condensin binding regions (Dowen *et al*, 2013). The fold-change values were normalized for each condition to the value of siCtrl cell samples. The normalized fold change values were represented in ESCs and MEFs on a scatter plot. More than three independent experiments were performed. FC, fold-change value.

## Discussion

We first studied how meiotic cohesins cooperate with mitotic cohesins to regulate sister chromatid cohesion and investigated their roles in chromosome organization, DNA replication, and repair during mitosis in ESCs. The meiotic cohesin components REC8 and STAG3 were specifically expressed in ESCs and were sufficient to function as essential cohesin components in ESCs. The relative contributions of the meiosis-specific cohesin to chromosome structure and cellular functions in mitotic programs have not been revealed. This study has led to several new findings which provide deeper insights into the mechanism that controls the morphogenesis of chromosomes and sister chromatid cohesion, which relies on a delicate balance between the mitotic and meiotic cohesin complexes in mitotic ESC chromosomes.

### REC8–STAG3 cohesin complexes support sister chromatid cohesion in ESC chromosome

REC8 and STAG3 are core cohesin factors that regulate sister chromatid cohesion in ESCs. REC8- or STAG3-null ESCs were found to have major defects in metaphase chromosome cohesion but were nevertheless capable of forming the metaphase plate with precociously separated sister chromatids. Interestingly, ESCs simultaneously expressed both REC8 and STAG3, and these proteins were copurified from nuclear fractions. This new finding supports the possibility that ESCs systematically express STAG3 to regulate the translocation of REC8 into the nucleus. Therefore, REC8-containing cohesin complexes in the nucleus could function with mitotic cohesin complexes and specifically mediate ESC-specific cohesin roles. These findings build upon those of a previous study that revealed that ectopically expressed-REC8 in Hek293 cells requires STAG3 to functionally replace RAD21, the mitotic counterpart of REC8 (Wolf *et al*, 2018).

### REC8-cohesin complexes support the centromeric cohesion in ESCs

It has been previously shown that centromeric cohesion is critical for precise chromosome segregation and is likely the reason for cohesin accumulation at centromeres during mitosis (Hirota *et al*, 2004). During the metaphase-to-anaphase transition, when all chromosomes have bioriented mitotic spindles, the cysteine protease separase (ESPL1/ESP1) cleaves the centromeric RAD21 protein to finally separate the sister chromatids (Hong *et al*, 2019; Hauf *et al*, 2001; Hearing & Nasmyth 2003; Uhlmann *et al*, 2000; Ciosk *et al*, 1998). In our study, REC8 but not RAD21 was abundantly localized in centromeres at metaphase, implying that REC8-mediated cohesion is maintained to hold sister chromatids at centromeres before anaphase onset. Thus, to suppress the precocious separation of sister chromatids, REC8-cohesin complexes might more effectively stabilize centromeric cohesion than RAD21-cohesin complexes. Since the separase effectively recognizes phosphorylated REC8 (Yoon *et al*, 2016; Kudo *et al*, 2009; Katis *et al*, 2010), a kinase or its regulatory factor may control its phosphorylation to avoid separation.

### Meiotic cohesin components modulate chromosome compaction in ESCs

Chromosomes have been shown to undergo global morphological changes between the compacted and expanded stages (Kleckner 2006). Mitotic chromosomes are compacted in preparation for the chromosome segregation that occurs during preanaphase and maintain a parallel alignment of sister chromatids (Liang *et al*, 2015; Batty & Gerlich 2019). The organization of chromosomes related to compaction progresses from prophase to metaphase via cycles of stress accumulation and release (Liang *et al*, 2015). A recent study using high-resolution 3D imaging suggested that “mini-axis” bridges containing cohesin link the split sister axes of mitotic chromosomes to provide mechanical stability during mitotic prophase (Chu *et al*, 2020) (Figure 8A). Collectively, these findings suggest that mechanical stress within the chromosome axes might modulate whole-chromosome morphology.

**Figure 8.**
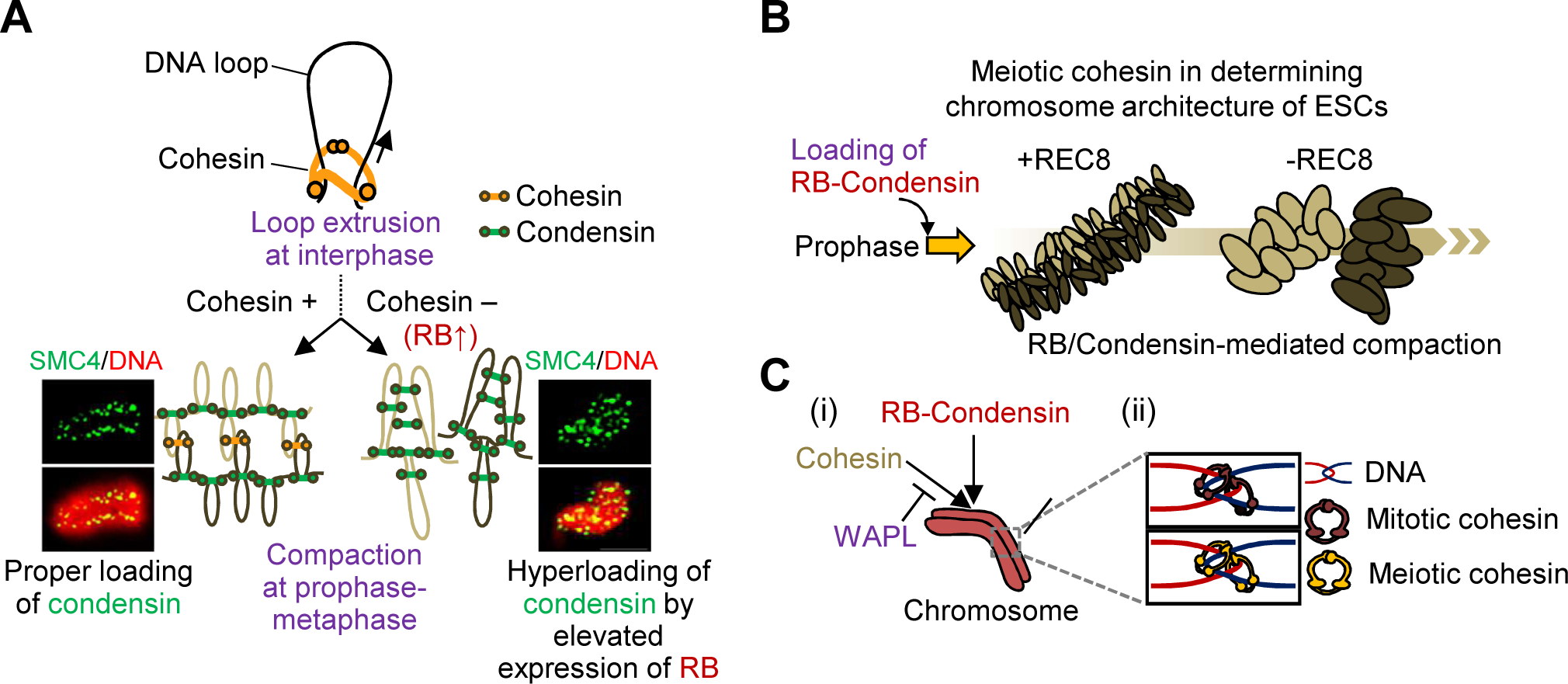
Meiotic cohesin components mediate sister chromatid cohesion and chromosomal organization in ESCs. (**A**) Regulation of chromosome structure by cohesin and condensin. (**B**) Proposed model of chromosome morphogenesis in normal or aberrant sister chromatid cohesion in ESCs. Loss of REC8-containing cohesin induces high level RB expression and enhances the compaction of chromosome structures from prophase to metaphase. (**C**) Cohesin/Condensin loading onto chromosomes. (i) The WAPL-mediated removal of cohesin recruits RB-condensin complexes and promotes chromosome condensation. In the absence of WAPL, cohesin removal is suppressed and condensin cannot be efficiently loaded on the chromosomes. (ii) Distinct localization of mitotic and meiotic cohesins at cohesion sites in ESCs.

Cohesin and condensin could be involved in the regulation of chromosome compaction during prophase in both mitosis and meiosis (Hong *et al*, 2019; Challa *et al*, 2016; Lazar-Stefanita *et al*, 2017). It has been shown in yeast meiosis that the absence of cohesin components induces the longitudinal hypercompaction of meiotic chromosomes during prophase (Hong *et al*, 2019). Condensins are localized along the central axes of the mitotic chromosomes to allow linkage between adjacent and/or distant DNA loops in prophase-to-metaphase transition (Figure 8A). In interphase, DNA loops are formed, extruded, and tethered together by cohesin (Abramo *et al*., 2019). In the absence of cohesin complexes, high levels of RB-condensin complexes bind widely to chromosomes, form new loops, and induce long-range looping by tethering adjacent loops. As the cells progress into metaphase, the absence of REC8- or RAD21-cohesin complexes causes precocious separation of sister chromatids and hyperloading of RB-condensin complexes on mitotic chromosomes that leads to hypercompaction of chromosomes (Fig 8A). Mitotic and meiotic cohesin complexes are loaded onto ESC chromosomes during the interphase, which is followed by cohesin dissociation and condensin loading in the early prophase. However, the absence of the REC8- cohesin complexes distributes condensin widely to mitotic chromosomes and creates short and fat chromosomes. This implies that meiotic cohesin is essential to modulate chromosome morphogenesis, even in the presence of the RAD21-cohesin complexex (Fig 8B). Thus, we propose that REC8/STAG3-cohesin complex is an essential factor for modulating chromosome morphogenesis in ESCs. Additional data suggest that chromosome loop size increases while axis length decreases progressively during the compaction process (Gibcus *et al*, 2018). General DNA loop extrusion during interphase is regulated by cohesins, but recent studies have suggested that condensin complexes can contribute to the same mechanism (Hirano 2016; Yamashita *et al*, 2011). However, we could not detect SMC4 protein in ESCs or MEFs during interphase, and the loss of cohesins showed that the condensins were randomly distributed to chromosome bodies during metaphase. Additionally, the pattern observed in prophase was similar in fixed whole nuclei and in chromosome spread samples, supporting that the condensins were bound all over the chromosome after cohesin knockdown, which caused hypercompaction. Thus, a timely association and/or dissociation of condensin and cohesin is important to promote normal chromosome compaction during mitosis in ESCs.

RNA Sequencing datasets with knockdown of cohesin components (SMC3, REC8, and RAD21) demonstrated that transcripts of RB were significantly increased in cohesin knockdown ESCs as compared with expression in control cells. Furthermore, RB actively interacts with condensin complex and mediates mitotic chromosome condensation (Coschi *et al*, 2014; this study). These imply that the high level expression of RB on aberrant sister chromatid cohesion enhances hyperloading of condensin on chromosomes that causes hypercompaction of chromosomes. We thus suggest that cohesin-mediated remodeling signals regulate chromosome compaction as a consequence of molecular redistribution. These findings offer a new framework for additional phenomena during chromosome compaction mediated by mitotic/meiotic cohesin and condensin.

### Cohesin and condensin compete for their binding sites

During the cell cycle, cohesin bound to interphase chromatin are released during prophase, coincident with their association with condensin (Abramo *et al*., 2019; Losada 2002). Generally, condensins participate in the mitotic phase of chromosome assembly in early prophase and promote chromosome compaction in preparation for mitotic chromosome segregation (Hagstrom & Meyer 2003; Hirota *et al*, 2004). However, the mechanism by which condensins achieve the enormous task of timely compaction of the entire genome remains mostly unknown. Chromosome compaction also requires cohesin, which support the close alignment of sister chromatids at an earlier point than condensin (Peters *et al*, 2008; Skibbens 2019; Guillou *et al*, 2010; Lazar-Steganita *et al*, 2017). Thus, our results suggested that both cohesin and condensin are involved in the modulation of chromosome structure and play similar roles in ensuring timely compaction. It has been previously shown that cohesin and condensin overlap at the same binding sites in the genome (Dowen *et al*, 2013; Kagey *et al*, 2010). These observations allowed us to propose a model for the cooccupation of cohesin/condensin binding sites—that is, condensin can gain greater access to chromosomes and attach to cohesin binding sites by cooccupying these binding sites when cohesin components are absent (Fig 8C). Our co-IP analysis identified that the level of condensin bound to cohesin binding sites was increased when we depleted several mitotic or meiotic cohesin components in ESCs. This indicates that condensin regulates the chromosome structure by directly binding to cohesin binding sites. Further, hyperloading of RB-condensin can be replaced with cohesins and are widely/irregularly distributed along prophase and metaphase chromosomes after knockdown of cohesin. However, given that cohesin binding is restricted to open chromatin sites, the possibility that condensin regulates chromosome compaction by binding at cohesin binding sites as well as other binding sites cannot be excluded.

### REC8–STAG3 cohesin complexes are required for the maintenance of genome integrity in ESCs

Chromosomes must undergo precise topological changes during diverse phases of the cell cycle. These processes must be executed with precision to avoid genome instability and whole-chromosome aneuploidy, which can cause tumorigenesis, cell death, and congenital disorders (Gruhn *et al*, 2019; Santaguida & Amon 2015; Ottolini *et al*, 2016). In addition to its function in sister chromatid cohesion, cohesin plays an important role in diverse endogenous and exogenous stresses (Hopkins *et al*, 2014; Mehta *et al*, 2013). Cohesin accumulates at DNA double-strand break (DSB) sites and is required for proper DNA repair, which relies on proper sister chromatid cohesion (Michaelis & Nasmyth 1997; Watrin & Peters 2006; Potts *et al*, 2006). Previous studies have demonstrated that the absence of Rec8 in budding yeast leads to a considerable decrease in the rate of the formation of recombinant products, suggesting its role in programmed DSB repair processes through recombination during meiosis (Hong *et al*, 2019; Kim *et al*, 2010; Mehta *et al*, 2013; Hong *et al*, 2013). Although the roles of REC8 in the regulation of chromosome segregation and genetic recombination have been reported during meiosis (Kim *et al*, 2010), there has been no definitive study on its role during the mitotic cell cycle. We found that lower expression of the REC8 or RAD21 genes resulted in a significant increase in the proportion of apoptotic cells in ESCs. Considering the data above, it is possible that the cohesin complexes composed of REC8 and RAD21 are functionally linked, and the meiotic cohesin REC8 may play an essential role in the maintenance of chromosome structural stability and cohesion-mediated cellular progression of ESCs. Thus, REC8/STAG3-cohesin complexes directly participate in faithful DNA replication and DNA repair pathways during mitosis. Consistent with this finding, we showed that the loss of REC8 and STAG3 delayed the progression of the replication fork during S phase and induced the accumulation of RAD51 and RPA foci at DNA break sites, indicating that the knockdown of REC8 led to large, unrepaired ssDNA gaps, ultimately affecting genome integrity.

## Conclusion

Based on the findings outlined above, we propose that REC8 and STAG3 are key components of the mitotic cohesin complex in ESCs. In addition to its general role for the organization of chromosome topological properties as seen in RAD21-cohesin complexes of ESCs, REC8-cohesin complexes specifically maintains centromeric cohesion. Furthermore, high resolution 3D imaging revealed that after knockdown of REC8- or RAD21-cohesin, hyperloading of RB-condensin complexes on chromosomes induces chromosome hypercompaction from prophase onward. These new insights show that how mitotic and meiotic cohesins are able to promote distinct chromosome interactions and that chromosome morphogenesis is regulated by RB-condensin interaction in the presence or absence of mitotic and meiotic cohesin factors. Our findings thus provide answers to the questions regarding chromosome compaction programs and sister chromatid cohesion in the mitotic ESC chromosomes.

## Materials and Methods

### Cell lines

Murine ESCs (J1) and MEFs were prepared as described previously (Choi *et al*, 2018). ESCs were cultured on 0.1% gelatin-coated 60-mm dishes in DMEM-high glucose (Gibco; 10569) supplemented with 10% horse serum (Gibco; 16050122), 10 mM HEPES (Gibco; 15630080), 2 mM L-glutamine (Gibco; 25030081), 0.1 mM Minimum Essential Medium Non-Essential Amino Acids (Gibco; 11140050), 0.1 mM β-mercaptoethanol (Gibco; 21985023), 100 U/ml Penicillin-Streptomycin (Gibco; 15140122), and 10^3^ U/mL ESGRO recombinant mouse leukemia inhibitory factor (LIF) (Millipore; ESG1107) in an humidified cell incubator with 5% CO_2_ at 37°C.

### Immunoprecipitation

Cell lysates from ESCs and MEFs were prepared as described previously (Choi *et al*, 2018). For the immunoprecipitation analysis, cells were cultured to a final cell concentration of 5x10^6^ per 60-mm culture dish. Cells were washed with PBS to remove the extra medium and treated 0.5% Trypsin-EDTA for 3 min at 37°C. Cells were then resuspended with PBS and centrifuged them at 1,000 rpm for 3 min. Cell pellets were resuspended with protein lysis buffer (50 mM Tris-HCl pH 7.6, 250 mM NaCl, 1 mM DTT, 5 mM MgCl_2_, 0.05% NP40, and 0.1 mM EDTA) for 10 min on ice. After centrifugation (13,500 rpm for 10 min), the protein lysates were transferred to microtubes and determined protein concentration using Bradford assay. Cell lysates (700 μg) were incubated with 1^st^ 1 μg antibodies overnight and mixed with 20 μl 50% protein-A/G agarose beads for 3 h at 4°C. Samples were centrifuged at 3000 rpm for 3 min at 4°C and the supernatants were discarded. Protein-A/G agarose beads were washed three times with washing buffer (50 mM HEPES, 2.5 mM MgCl_2,_ 100 mM KCl, 10% glycerol, 1 mM DTT, and 0.1% NP40 with 1x PIC), and PBS. After centrifugation (3000 rpm for 3 min at 4°C), agarose beads were mixed with 20 μl 2⍰ concentrated SDS sample buffer and boiled for 5 min at 100°C. Then, protein samples were transferred to fresh microtubes and gel electrophoresis was carried out.

### Mouse spermatocyte immunofluorescent staining

Male spermatocyte cells were taken from 4 weeks old testes of C57/BL6J strain purchased from Jackson laboratory (Bar Harbor, Maine, USA) via Orient Bio (Seongnam, Korea). Meiotic spread preparation and immunofluorescent staining method were prepared as described (Yoon *et al*, 2018).

### Antibodies

The source of the antibodies are as follows: anti-rabbit SMC3 (Abcam; ab9263; 1:5,000), anti-rabbit RAD21 (Abcam; ab154769; 1:5,000), anti-rabbit STAG3 (Abcam; ab185109; 1:1,000), anti-rabbit REC8 (Abcam; ab192241; 1:3,000), anti-rabbit SMC4 (Abcam; ab17958; 1:5,000), anti-rabbit α−tubulin (Abcam; ab4074; 1:10,000), anti-mouse IdU (BD; B44; 1:25), anti-rat BrdU (Abcam; ab6326; 1:500), anti-mouse OCT3/4 (Santa Cruz; sc-5297; 1:3,000), anti-rat RPA (Cell signaling; 2208; 1:3,000), anti-mouse RAD51 (Merck Millipore; PC130; 1:3,000), anti-rabbit RB (Abcam; ab181616; 1:2000), anti-mouse RB (Abcam; ab24; 1:2000) and anti-mouse SYCP3 (Santa Cruz; sc- 74569; gift from National Cancer Center in Korea; 1:500). We used the following secondary antibodies: anti-mouse Alexa 488 (111-545-003, 1:500; Jackson), anti-rat fluorescein isothiocyanate (FITC) (112- 095-003, 1:500; Jackson), anti-rabbit TRITC (115-025-003), and anti-rabbit Cy3 (111- 165-003, 1:500; Jackson). We have produced a rabbit polyclonal antibody against the recombinant C-terminal 342 amino acid sequences of REC8 (GenBank accession numbers: mouse REC8, BC052155.1).

### Probes for Fluorescence In Situ Hybridization

Telomere probes were prepared with the mouse repeat sequence (TTAGGG)_n_ and labeled it with 5’- FAM and 3’-FAM primers using a PCR reaction: 5’-6FAM- TTAGGGTTAGGGTTAGGGTTAGGGTTAGGG-3’ and 5’-6FAM-CCCTAACCCTAACCCTAACCCTAACCCTAA-3’. PCR reaction was performed in 50 μl containing 10 mM Tris-HCl (pH 8.3), 50 mM KCl, 1.5 mM MgCl_2_, 200 mM of each dNTP, 0.2 mM of each primer, and 2 units of Taq polymerase. The amplification was composed of 10 cycles of 1 min at 95°C, 30 sec at 54°C, and 60 sec at 72°C, followed by 30 cycles of 1 min at 95°C, 30 sec at 62°C, 90 sec at 72°C, and one final step of 7 min at 72°C. The locus-specific probe was bound to the RAD54 locus region (Chromosome 4: 116,094,264 – 116,123,690). Probes were labeled with Cy3- dCTP in the same PCR reaction. The PCR was carried out with mouse cDNA using the following primers: 5’-AGCACAGTGGCGGCCGCATGAGGAGGAGCTTA-3’ and 5’-TAGACTCGAGCGGCCGCTCAGTGAAGCCGCGC-3’. The BAC DNA was extracted with the NucleoBond Xtra Midi kit according to the manufacturer’s protocol. Purified BAC DNA were then labeled by standard nick translation reaction, including dATPs, dGTPs, dTTPs, Cy3-labeled dCTPs, translation buffer, diluted DNase I, DNA polymerase I, and BAC DNA. Mixture was incubated at 15°C for 2 h. To inactivate the reaction, stop solution (0.5 M EDTA and 10% SDS) was added and incubated at 68°C for 15 mins. Products were then prepared on a 1.2% agarose gel for the presence of a smear between 500 and 200 bp. Probes were stored at –20°C.

### Fluorescence in situ hybridization on metaphase chromosomes

For mitotic chromosome spreads, ESCs and MEFs were incubated with 0.1 μg colcemid (Gibco; 15212012) in 4 ml medium for 3 h to arrest the cells in the metaphase stage. Cells were resuspended in 75 mM KCl for 10 min at 37°C, and then resuspended the fixed cell pellets in methanol/acetic acid (3:1 v/v) and dropped them on slide glasses. Slide glasses were left overnight at room temperature. The chromosome preparation was rehydrated in 2X concentrated SCC buffer, digested with pepsin (1 mg/ml) in 10 mM HCl (pH 2.0) for 10 min at 37°C, fixed in 3.7% formaldehyde in PBS for 5 min and then washed with PBS twice for 5 min each time. The samples were rehydrated with series of incubation in 70%, 90%, and 100% cold ethanol and let them air dry. Slides were incubated with a pre-warmed (75°C) denaturation buffer for 2 min on the area of the slide marked with good metaphase chromosomes. For the hybridization step, both slides and the Cy3-labeled probe were pre-warmed at 37°C for 5 min and then 8 μl of hybridization mixture was placed on the spotted area of the slice for 1 h at 37°C. Slides were washed once in PBS, once in 2☓-concentration SCC buffer at room temperature for 10 min, and twice in PBS for 5 min each.

Slides were then mounted with antifade solution containing 4′,6-diamidino-2-phenylindole-dihydrochloride (1 μg/ml DAPI). To distinguish the cell cycle phases, bromodeoxyuridine (BrdU) (Sigma Corp.; B5002) was added to the growth medium and cells were cultured in a humidified incubator with 5% CO_2_ at 37°C for 15 min. Then, cells were harvested for the FISH analysis (Figure 1E). We measured the lengths of chromosomes corresponding to telomere- and locus-specific labeling using Nikon NIS software.

### Metaphase Spread Immunostaining

For the preparation of mitotic chromosome spreads, ESCs were treated with 0.1 μg/ml colcemid for 3 h in a humidified incubator with 5% CO_2_ at 37°C. The cells were hypotonically swelled with a hypotonic buffer (75 mM KCl, 0.8% Na Citrate, and H_2_O_2_ 1:1:1 v/v/v) for 10 min at 37°C. Three-volume of cold methanol-acetic acid (3:1) was added to the hypotonic buffer for 3 min. Supernatant was removed with an aspirator and 5 ml of cold methanol-acetic acid (3:1) was added to cell pellets and incubated for 5 min. We gently resuspended the cell pellets and put a drop on a slide. As soon as the surface of the slide was dry, the slides were rehydrated by dipping them in PBS-azide buffer (10 mM NaPO_4_ pH7.4, 0.15 M NaCl, 1 mM EGTA, and 0.01% NaN_3_) for 10 min. The slide was then swollen by washing the slides three times (3 min each) with TEEN buffer (1 mM Tris-HCl pH8.0, 0.2 mM EDTA, 25 mM NaCl, 0.5% Triton X-100, and 0.1% BSA). Primary antibody was then added on the slide for 1 h at room temperature. Primary antibody (diluted in TEEN) was removed by washing three times with KB washing buffer (10 mM Tris-HCl pH 8.0, 0.15 M NaCl, and 0.1% BSA) for 3 min. Fluorescein-conjugated secondary antibody (diluted in TEEN) was incubated on the slide for 1 h at room temperature. After a final washing with KB buffer, slides were mounted with antifade solution including DAPI. The metaphase cell images were captured using a fluorescence microscope (Nikon Eclipse, Ti-E, Japan).

### Immunostaining

ESCs and MEFs were cultured on poly-L-lysine-coated (Santa Cruz; sc-286689) 12 mm coverslips (Deckglaser; 0111520) and then fixed with 4% paraformaldehyde for 10 min. Cells were then permeabilized with 0.2% Triton X-100 for 5 min and washed with PBS-T (0.1% Tween 20 in PBS) for 3 min three times. Cells were blocked with 3% BSA in PBS-T for 30 min and then incubated for 1 h with the following primary antibodies (diluted in 1% BSA in PBS-T): anti-RPA (Cell Signaling, #2208, 1:200), anti-Rad51 (Santa Cruz, sc-8349, 1:200), anti-Rad54 (Santa Cruz, sc-374598, 1:200), and anti-Exo1 (Thermo, MS-1534-P, 1:100). After washing three times with T-PBS, cells were incubated for 1 h with the appropriate anti-Alexa488 (Jackson, 111-545-003, 1:500), anti-Fluorescein isothiocyanate (FITC) (Jackson, 112-095-003, 1:500), anti-Cy3 (Jackson, 111-165-003, 1:500) - conjugated secondary antibodies and then mounted with antifade solution containing DAPI. Images were captured using an Eclipse Ti-E fluorescence microscope (Nikon, Tokyo, Japan) equipped with fluorescence filters for DAPI and the indicated fluorophores conjugated to the secondary antibodies.

### Chromatin immunoprecipitation

For the chromatin immunoprecipitation assay, we cultured 4 x 10^5^ – 5 x 10^5^ ESCs and MEFs. Cells were chemically with 1% formaldehyde for 15 min at room temperature and added glycine (125 mM) for 5 min at room temperature. Cells were then lysed with SDS-lysis buffer (50 mM HEPES, 140 mM NaCl, 1 mM EDTA, 1% Triton X-100, 0.1% Sodium deoxycholate, 0.1% SDS, and 1⍰ Protease inhibitor) and solubilized by sonication. Cell lysates were incubated overnight at 4°C with 1 μg antibodies and added 20 μl protein A/G-agarose beads (GenDEPOT; P9203-050). For the ChIP analysis of SMC3, RAD21, REC8, and SMC4 in MEFs and ESCs, protein A/G-agarose beads were washed four times with a low-salt buffer (0.1% SDS, 1% Triton X-100, 2 mM EDTA, 20 mM Tris-HCl pH 8.0, and 150 mM NaCl), a high salt buffer (0.1% SDS, 1% Triton X-100, 2 mM EDTA, 20 mM Tris-HCl pH 8.0, and 500 mM NaCl), a LiCl buffer (0.25 M LiCl, 1% NP-40, 1% Na_2_ Deoxycholate, 1 mM EDTA, and 10 mM Tris-HCl pH 8.0), and a TE buffer (10 mM Tris pH 8.0, and 1 mM EDTA). The bound complexes were eluted from the beads with an elution buffer and reversed the crosslink with incubation at 65°C for 12 hr. The purified DNA was quantified with qPCR analysis.

### Quantitative PCR

To analyze the chromatin immunoprecipitations, purified DNA was quantified by qPCR using SYBR Green (Bioneer; K-6251) and the CFX Connect Real-Time PCR system (Bio-Rad; 1855201). We confirmed a subset of strongly bound cohesin sites overlapping with the binding sites of condensin (Dowen *et al*, 2013). The primer sequences used in quantitative PCR were as follows: Forward: 5’- GTGCTGGGATTAAAGGCG-3’ and reverse: 5’-AGATGAGGCTTTCAGGAAATAC-3’. For figure 4G, the primer sequences were as follows: (Major satellite) Forward: 5’- GACGACTTGAAAAATGACGAAATC-3’ and reverse: 5’-CATATTCCAGGTCCTTCAGTGTGC-3’ and (Minor satellite) Forward: 5’-CATGGAAAATGATAAAAACC-3’ and reverse: 5’- CATCTAATATGTTCTACAGTGTGG-3’.

### DNA fiber assay

ESCs and MEFs (2x10^5^ cells) were pulsed with 50 μM ldU (Glentham; GP1769) for 15 min and then sequentially pulsed with 100 μM CldU (Sigma; C6891) for 15 min. Cells were resuspended in cold PBS at a concentration of 400 cells per μl. Following the addition of 9 μl DNA lysis buffer (200 mM Tris-HCl pH 7.4, 0.5% SDS, and 50 mM EDTA), a drop of resuspended cells was put on a slide. Slides were tilted approximately 10° to allow the lysed DNA to spread and let it dry for 30 min. Cells were fixed with methanol-acetic acid (3:1 v/v) for 10 min and dry overnight at room temperature.

After washing with PBS, the slide was immersed in 2.5 N HCl for 60 min at room temperature to denature the DNA molecules. IdU incorporation was detected using a mouse anti-IdU antibody (BD; B44, 1:25) and BrdU incorporation was detected using a rat anti-BrdU antibody (Abcam; ab6326, 1:500). Slides were then washed with PBS and incubated with fluorescence secondary antibodies, TRITC-conjugated goat anti-mouse IgG antibody (Jackson lab; 115-025-003, 1:300) and FITC-labelled goat anti-rat IgG antibody (Jackson lab, 112-095-003, 1:400) to detect IdU and CldU, respectively. After the final wash with PBS, slides were mounted with antifade solution containing DAPI. DNA fiber images were captured using a fluorescence microscope (Nikon Eclipse, Ti-E, Japan) and measured their lengths according to the IdU and CldU labeling using the Nikon NIS software.

### RNA interference

We used a commercially available predesigned siRNA (Bioneer; AccuTarget^tm^) specific to the target genes to knockdown their endogenous expression in ESCs and MEFs. The siRNA pool included single targeting sequences as follows: SMC3: 5’-GAGGUUGGCUCAAGCUACA(dTdT)-3’, RAD21: 5’-GUGCAAUAUUGGUGCAUGU(dTdT)-3’, REC8: 5’-GAGAUCAGUCGAGGAGACU(dTdT)-3’, STAG3: 5’-CUGGAUUAACAUGCCUACU(dTdT)-3’ WAPL: 5’- GUCCUUGAAGAUAUACCAA(dTdT)-3’ Oligonucleotides were transfected using Lipofectamine RNAiMAX (Thermo Fisher; 13778150) according to the manufacturer’s instructions. We also used a negative control siRNA purchased from Bioneer (SN-1003). The siRNAs were treated in an optimum (serum-free) medium for 48 h.

### RNA-Seq library preparation and sequencing

Total RNA samples were isolated using the RNeasy Mini Kit (Qiagen). All the detailed applications were followed by the manufacturer’s direction. RNA quality was assessed by Agilent 2100 bioanalyzer (Agilent Technologies, Amstelveen, The Netherlands), and RNA concentration was determined using ND-2000 Spectrophotometer (Thermo Inc., DE, USA). mRNA-Seq libraries were prepared from total RNA using the NEBNext Ultra II Directional RNA-Seq Kit (NEW ENGLAND BioLabs, Inc., UK). The isolation of mRNA was performed using the Poly(A) RNA Selection Kit (LEXOGEN, Inc., Austria). The isolated mRNAs were used for the cDNA synthesis and shearing, following manufacture’s instruction. Indexing was performed using the Illumina indexes 1-12. The enrichment step was carried out using PCR reaction. Subsequently, libraries were checked using the Agilent 2100 bioanalyzer (DNA High Sensitivity Kit) to evaluate the mean fragment size. Quantification was performed using the library quantification kit using a StepOne Real-Time PCR System (Life Technologies, Inc., USA). High-throughput sequencing was performed as paired-end 100 sequencing using HiSeq X10 (Illumina, Inc., USA).

### Data analysis

A quality control of raw sequencing data was performed using the FastQC tool. Adapter and low-quality reads (< Q20) were removed using FASTX_Trimmer and BBMap software tools. Then the trimmed reads were mapped to the reference genome using TopHat. Gene expression levels were estimated using FPKM (Fragments Per kb per Million reads) values by Cufflinks. The FPKM values were normalized based on Quantile normalization method using edgeR in R. Data mining and graphic visualization were performed using ExDEGA software (E-biogen, Inc., Korea).

### Statistical analysis

Data were analyzed with GraphPad Prism 5 and OriginPro 2020b software. The data were described as the means ± SD. We measured the statistically significant differences between various groups using unpaired student’s t-tests. The statistical significance was set at *p < 0.05, **p < 0.01, and ***p < 0.001. In this study, we conducted and analyzed more than three independent experiments. For Figures 2B and E, we used box and whisker plots to analyze the distribution of the

DNA fiber lengths. Top and bottom dots mean high and low rank 10% respectively, while the box plot indicates the lower quartile. Seventy-five percent of the scores fell below the upper quartile value, the median, which averages the values of all data, and the low quartile, which twenty-five percent of scores fell below. We performed t-tests (Mann–Whitney) using the GraphPad Prism 5 software (ns: not-significant, ***P < 0.001). For Figures 2C and F, the DNA fiber lengths were converted to kilobase (kb) (1 mm = 2.94 kb) to calculate the DNA fork rates and the converted values divided by CldU/IdU pulse labeling time (40 min). P-values were Student’s t-test (ns: not- significant, ***P < 0.001, and n, the number of measured fibers). For Figures 4B, E and I, error bars represent mean ± SD values of the three independent experiments and counted at least 30 chromosomes. P-values (Student’s t-test) were calculated using GraphPad Prism 5 software. For Figures 5 E-F and J-K, a violin box plot was used to analyze the distribution of condensin. The box whisker shows the five-number as a dataset, including the minimum score, lower quartile, median, upper quartile, and maximum score. P-values (Student’s t-test) were calculated using OriginPro 2020b (ns: not-significant, ***P < 0.001).

### Data availability

RNA Sequencing data have been deposited in the NCBI Sequence Read Archive. All RNA-Seq reads are available under the following accession numbers: SRX10686134 (https://www.ncbi.nlm.nih.gov/sra/?term=SRX10686134), SRX10686135 (https://www.ncbi.nlm.nih.gov/sra/?term=SRX10686135), SRX10686136 (https://www.ncbi.nlm.nih.gov/sra/?term=SRX10686136), and SRX10686137 (https://www.ncbi.nlm.nih.gov/sra/?term=SRX10686137).

## Acknowledgements

We thank the microscopy core facility of SNU medical school for the help and expertise with running 3D SIM. This work was supported by grants to K.P.K. from the National Research Foundation of Korea, funded by the Ministry of Science, ICT & Future Planning (No. 2020R1A2C2011887; 2018R1A5A1025077) and the Next-Generation BioGreen 21 Program (No. PJ015708), Rural Development Administration, Republic of Korea.

## Author contribution

E.H.C., Y.E.K, and S.B.Y. designed the experiments and performed most the experiments. K.P.K. conceived and supervised the project. E.H.C., Y.E.K., Y.H., and K.P.K. analyzed the results and prepared the manuscript, with input from all authors.

## References

1. Abramo K, Valton A-L, Venev SV, Ozadam H, Fox AN, Dekker J (2019) A chromosome folding intermediate at the condensin-to-cohesin transition during telophase. Nat. Cell. Biol. 21: 1393–1402

2. Ahuja AK, Jodkowska K, Teloni F, Bizard AH, Zellweger R, Herrador R, Ortega S, Hickson ID, Altmeyer M, Mendez J et al (2016) A short G1 phase imposes constitutive replication stress and fork remodelling in mouse embryonic stem cells. Nat. Commun. 7: 10660

3. Batty P, Gerlich DW (2019) Mitotic chromosome mechanics: How cells segregate their genome. Trends in Cell Biol. 29: 717–726

4. Biswas U, Hempel K, Llano E, Pendas A, Jessberger R (2016) Distinct roles of meiosis-specific cohesin complexes in mammalian spermatogenesis. PLos Genet. 12: e1006389

5. Brar GA, Hochwagen, A, Ee LS, Amon A (2009) The multiple roles of cohesin in meiotic chromosome morphogenesis and pairing. Mol. Biol. Cell. 20: 1030–1047

6. Carretero M, Ruiz-Torres M, Rodríguez-Corsino M, Barthelemy I, Losada A (2013) Pds5B is required for cohesion establishment and Aurora B accumulation at centromeres. EMBO J. 32: 2938–2949

7. Challa K, Lee MS, Shinohara M, Kim KP (2016) Shinohara A. Rad61/Wpl1 (Wapl), a cohesin regulator, controls chromosome compaction during meiosis. Nucleic Acids Res. 44: 3190–3203

8. Choi EH, Yoon S, Park KS, Kim KP (2017) The homologous recombination machinery orchestrates post-replication DNA repair during self-renewal of mouse embryonic stem cells. Sci. Rep. 7: 11610

9. Choi EH, Yoon S, Kim KP (2018) Combined ectopic expression of homologous recombination factors promotes embryonic stem cell differentiation. Mol. Ther. 26: 1154–1165

10. Choi EH, Yoon S, Koh YE, Kim KP (2020) Maintenance of genome integrity and active homologous recombination in embryonic stem cells. Exp. Mol. Med. 52: 1220–1229

11. Chu L, Liang Z, Mukhina M, Fisher J, Vincenten N, Zhang Z, Hutchinson J, Zickler D, Kleckner N (2020) The 3D topography of mitotic chromosomes. Mol. Cell 79: 902–916

12. Ciosk R, Zachariae W, Michaelis C, Shevchenko A, Mann M, Nasmyth K (1998) An Esp1/Pds1 complex regulates loss of sister chromatid cohesion at the metaphase to anaphase transition in yeast. Cell 93: 1067–1076

13. Cockram C, Thierry A, Gorlas A, Lestini R, Koszul R (2020) Euryarchaeal genomes are folded into SMC-dependent loops and domains, but lack transcription-mediated compartmentalization. Mol. Cell 22: 30905–30909

14. Coelho PA, Queiroz-Machdo J, Sunkel CE (2003) Condensin-dependent localisation of topoisomerase II to an axial chromosomal structure is required for sister chromatid resolution during mitosis. J. Cell Sci. 116: 4763–4776

15. Coschi CH, Martens AL, Ritchie K, Francis SM, Chakrabarti S, Berube NG, and Dick FA (2010). Mitotic chromosome condensation mediated by the retinoblastoma protein is tumor-suppressive. Genes Dev. 24: 1351–1363

16. Coschi CH, Ishak CA, Gallo D, Marshall A, Talluri S, Wang J, Cecchini MJ, Martens AL, Percy V, Welch I et al (2014) Haploinsufficiency of an RB-E2F1-Condensin II complex leads to aberrant replication and aneuploidy. Cancer Discovery 4: 840–853

17. Díaz-Martínez LA, Giménez-Abián JF, Clarke DJ (2007) Cohesin is dispensable for centromere cohesion in human cells. Plos one 2: e318

18. Dixon JR, Jung I, Selvaraj S, Shen Y, Antosiewicz-Bourget J E, Lee AY, Ye Z, Kim A, Rajagopal N, Xie W et al (2015) Chromatin architecture reorganization during stem cell differentiation. Nature 518: 331–336

19. Dowen JM, Bilodeau S, Orlando DA, Hübner MR, Abraham BJ, Spector DL, Young RA (2013) Multiple structural maintenance of chromosome complexes at transcriptional regulatory elements. Stem Cell Reports 1: 371–378

20. Fukuda T, Fukuda N, Agostinho A, Hernández-Hernández A, Kouznetsova A, Höög C (2014) STAG3-mediated stabilization of REC8 cohesin complexes promotes chromosome synapsis during meiosis. EMBO J. 33: 1243–1255

21. Gandhi R, Gillespie P, Hirano T (2006) Human Wapl is a cohesin-binding protein that promotes sister-chromatid resolution in mitotic prophase. Curr. Biol. 16: 2406–2417

22. Gibcus JH, Samejima K, Goloborodko A, Samejima I, Naumova N, Nuebler J, Kanemaki MT, Xie L, Paulson JR, Earnshaw WC (2018) A pathway for mitotic chromosome formation. Science 359: eaao6135

23. Giménez-Abián JF, Sumara I, Hirota T, Hauf S, Gerlich D, de la Torre C, Ellenberg J, Peters JM (2004) Regulation of sister chromatid cohesion between chromosome arms. Curr. Biol. 14: 1187–1193

24. Goloborodko A, Imakaev MV, Marko JF, Mirny L (2016) Compaction and segregation of sister chromatids via active loop extrusion. Elife 18: e14864

25. Gruber S, Hearing CH, Nasmyth K (2003) Chromosomal cohesin forms a ring. Cell 112: 765–777

26. Gruhn JR, Zielinska AP, Shukla V, Blanshard R, Capalbo A, Cimadomo D, Nikiforov D, Chan AC et al (2019) Chromosome errors in human eggs shape natural fertility over reproductive life span. Science 365: 1466–1469

27. Guillou E, Ibarra A, Coulon V, Casado-Vela J, Rico D, Casal I, Schwob E, Losada A, Méndez J (2010) Cohesin organizes chromatin loops at DNA replication factors. Genes & Dev. 24: 2812–2822

28. Haarhuis JH, van der Weide RH, Blomen VA, Yáñez-Cuna JO, Amendola M, van Ruiten MS, Krijger P, Teunissen H, Medema RH, van Steensel B (2017) The cohesin release factor WAPL restricts chromatin loop extension. Cell 169: 693–707

29. Haarhuis JH, Elbatsh AM, Rowland BD (2014) Cohesin and its regulation: on the logic of X- shaped chromosomes. Dev. Cell 31: 7–18

30. Hagstrom KA., Meyer BJ (2003) Condensin and cohesin: more than chromosome compactor and glue. Nat. Rev. Genet. 4: 520–534

31. Hauf S, Waizenegger IC, Peters JM (2001) Cohesin cleavage by separase required for anaphase and cytokinesis in human cells. Science 293: 1320–1323

32. Hearing CH, Nasmyth K (2003) Building and breaking bridges between sister chromatids. Bioessays 25: 1178–1191

33. Hirano T (2016) Condensin-based chromosome organization from bacteria to vertebrates. Cell 164: 847–857

34. Hirano T, Kobayashi R, Hirano M (1997) Condensins, chromosome condensation protein complexes containing XCAP-C, XCAP-E and a Xenopus homolog of the Drosophila barren protein. Cell 89: 511–521

35. Hirota T, Gerlich D, Koch B, Peters JM (2004) Distinct functions of condensin I and II in mitotic chromosome assembly. J. Cell Sci. 117: 6435–6445

36. Hopkins J, Hwang G, Jacob J, Sapp N, Bedigian R, Oka K, Overbeek P, Murray S, Jordan PW (2014) Meiosis-specific cohesin component, Stag3 is essential for maintaining centromere chromatid cohesion, and required for DNA repair and synapsis between homologous chromosomes. PLoS Genet. 10: e1004413

37. Hong S, Sung Y, Yu M, Lee MS, Kleckner N, Kim KP (2013) The logic and mechanism of homologous recombination partner choice. Mol. Cell 51: 440–453

38. Hong S, Joo JH, Yun H, Kleckner N, Kim KP (2019) Recruitment of Rec8, Pds5 and Rad61/Wapl to meiotic homolog pairing, recombination, axis formation and S-phase. Nucleic Acids Res. 47: 11691–11708

39. Kagey MH, Newman JJ, Bilodeau S, Zhan Y, Orlando DA, van Berkum NL, Ebmeier CC, Goossens J, Rahl PB, Levine SS et al (2010) Mediator and cohesin connect gene expression and chromatin architecture. Nature 467: 430–435

40. Katis VL, Lipp JJ, Imre R, Bogdanova A, Okaz E, Habermann B, Mechtler K, Nasmyth K, Zachariae W (2010) Rec8 rhosphorylation by casein kinase 1 and Cdc7-Dbf4 kinase regulates cohesin cleavage by separase during meiosis. *Dev*, Cell 18: 397–409

41. Kim KP, Weiner BM, Zhang L, Jordan A, Dekker J, Kleckner N (2010) Sister cohesion and structural axis components mediate homolog bias of meiotic recombination. Cell 143: 924–937

42. Kleckner N (2006) Chiasma formation: chromatin/axis interplay and the role(s) of the synaptonemal complex. Chromosoma. 115: 175–194

43. Kschonsak M, Hearing CH (2015) Shaping mitotic chromosomes: From classical concepts to molecular mechanisms. Bioessays 37: 755–766

44. Kudo NR, Anger M, Peters A, Stemmann O, Theussl HC, Helmhart W, Kudo H, Heyting C, Nasmyth K (2009) Role of cleavage by separase of the Rec8 kleisin subunit of cohesin during mammalian meiosis I. J. Cell Sci. 122, 2686–2698

45. Lazar-Stefanita L, Scolari VF, Mercy G, Muller H, Guérin TM, Thierry A, Mozziconacci J, Koszul R (2017) Cohesins and condensins orchestrate the 4D dynamics of yeast chromosomes during the cell cycle. EMBO J. 36: 2684–2697

46. Lee MS, Joo JH, Kim KP (2017) Roles of budding yeast Hrr25 in recombination and sporulation. Journal of Microbiology and Biotechnology 27: 1198–1203

47. Liang Z, Zickler D, Prentiss M, Chang FS, Witz G, Maeshima K, Kleckner N (2015) Chromosomes progress to metaphase in multiple discrete steps via global compaction/expansion cycles. Cell 161: 1124–1137

48. Litwin I, Pilarczyk E, Wysocki R (2018) The emerging role of cohesin in the DNA damage response. Genes 9: 581

49. Liu L, Michowski W, Kolodziejczyk A, Sicinski P (2019) The cell cycle in stem cell proliferation, pluripotency and differentiation. Nat. Cell Biol. 21: 1060–1067

50. Losada A (2002) Cohesin release is required for sister chromatid resolution, but not for condensin-mediated compaction, at the onset of mitosis. Genes Dev. 16: 3004–3016

51. Maia AG, Alajem A, Meshorer E, Santos MR (2011) Open chromatin in pluripotency and reprogramming. Nat. Rev. Mol. Cell Biol. 12: 36–47

52. Mehta GD, Kumar R, Srivastava S, Ghosh SK (2013) Cohesin: Functions beyond sister chromatid cohesion. FEBS Lett. 587: 2299–2312

53. Michaelis C, Nasmyth K (1997) Cohesins: chromosomal proteins that prevent premature separation of sister chromatids. Cell 91: 35–45

54. Nasmyth K, Hearing CH (2009) Cohesin: Its roles and mechanisms. Annu. Rev. Genet. 43: 525–558

55. Ottolini CS, Newnham L, Capalbo A, Natesan SA, Joshi HA, Cimadomo D, Griffin DK, Sage K, Summers MC, Thornhill AR et al (2016) Genome-wide recombination and chromosome segregation in human oocytes and embryos reveal selection for maternal recombination rates. Nat. Genet. 47: 727–735

56. Onn I, Heidinger-Pauli JM, Guacci V, Unal E, Koshland DE (2008) Sister chromatid cohesion: a simple concept with a complex reality. Annu. Rev. Cell Dev. Biol. 24: 105–129

57. Peters JM, Tedeschi A, Schmitz J (2008) The cohesin complex and its roles in chromosome biology. Genes & Dev. 22: 3089–3114

58. Peters JM, Nishiyama T (2012) Sister Chromatid Cohesion. Cold Spring Harb. Perspect. Biol. 4: a011130

59. Potts P, Porteus MH, Yu H (2006) Human SMC5/6 complex promotes sister chromatid homologous recombination by recruiting the SMC1/3 cohesin complex to double-strand breaks. EMBO. J. 25: 3377–3388

60. Santaguida S, Amon A (2015) Short- and long-term effects of chromosome mis-segregation and aneuploidy. Nat. Rev. Mol. Cell Biol. 16: 473–485

61. Schöckel L, Möckel M, Mayer B, Boos D, Stemmann O (2011) Cleavage of cohesin rings coordinates the separation of centrioles and chromatids. Nat. Cell Biol. 13: 966–972

62. Sherwood R, Takahashi TS, Jallepalli PV (2010) Sister acts: coordinating DNA replication and cohesion establishment. Genes & Dev. 24: 2723–2731

63. Skibbens RV (2019) Condensins and cohesins – one of these things is not like the other! *J*. Cell Sci. 132: jcs.220491

64. Tee WW, Reinberg D (2014) Chromatin features and the epigenetic regulation of pluripotency states in ESCs. Development 141: 2376–2390

65. Uhlmann F, Wernic D, Poupart MA, Koonin EV, Nasmyth K (2000) Cleavage of cohesin by the CD clan protease separin triggers anaphase in yeast. Cell 103: 375–386

66. Vitale I, Manic G, De Maria R, Kroemer G, Galluzzi L (2017) DNA damage in stem cells. Mol. Cell 66: 306–319

67. Waizenegger IC, Hauf S, Meinke A, Peters JM (2000) Two Distinct Pathways Remove Mammalian Cohesin from Chromosome Arms in Prophase and from Centromeres in Anaphase. Cell 103: 399–410

68. Watrin E, Peters JM (2006) Cohesin and DNA damage repair. Exp. Cell Res. 312: 2687–2693

69. Watrin E, Peters JM (2009) The cohesin complex is required for the DNA damage-induced G2/M checkpoint in mammalian cells. EMBO J. 28: 2625–2635

70. Weitzer S, Uhlmann F (2002) Chromosome segregation: playing polo in prophase. Dev. Cell 2: 381–382

71. Wolf PG, Ramos AC, Kenzel J, Neumann B, Stemmann O (2018) Studying meiotic cohesion in somatic cells reveals that Rec8-containing cohesion requires Stag3 to function and is regulated by Wapl and sororin. J. Cell Sci. 131: jcs212100

72. Wu N, Yu H (2012) The Smc complexes in DNA damage response. Cell Biosci. 2: 5

73. Yamashita D, Shintomi K, Ono T, Gavvovidis I, Schindler D, Neitzel H, Trimborn M, Hirano T (2011) MCPH1 regulates chromosome condensation and shaping as a composite modulator of condensin II. J. Cell Biol. 194: 841–854

74. Yoon SW, Lee MS, Xaver M, Zhang L, Hong S, Kong J, Cho H, Kleckner N, Kim KP (2016) Meiotic prophase roles of Rec8 in crossover recombination and chromosome structure. Nucleic Acids Res. 44: 9296–9314

75. Yoon S, Choi EH, Kim J, Kim KP (2018) Structured illumination microscopy imaging reveals localization of replication protein A between chromosome lateral elements during mammalian meiosis. Exp. Mol. Med. 50: 10.1038/s12276-018-0139-5

76. Yu HG, Koshland D (2005) Chromosome morphogenesis: condensin-dependent cohesin removal during meiosis. Cell 123: 397–407

77. Zhang N, Kuznetsov SG, Sharan SK, Li K, Rao PH, Pati DA (2008) A handcuff model for the cohesin complex. J. Cell Biol. 183: 1019–1031

78. Zhang N, Kuznetsov SG, Sharan SK, Li K, Rao PH, Pati DA (2006) Wapl controls the dynamic association of cohesin with chromatin. Cell 127: 955–967

79. Zielinska AP, Bellou E, Sharma N, Frombach A, Seres KB, Gruhn JR, Blayney M, Eckel H, Moltrecht R, Elder K et al (2019) Meiotic kinetochores fragment into multiple lobes upon cohesin loss in aging eggs. Curr. Biol. 29: 3749–3765

80. Zwaka TP, Thomson JA (2003) Homologous recombination in human embryonic stem cells. Nat. Biotechnol. 21: 319–321

