## Supplemental Figures and Tables for "Meiosis-Specific Cohesin Complexes Display Distinct and Essential Roles in Mitotic ESC Chromosomes"

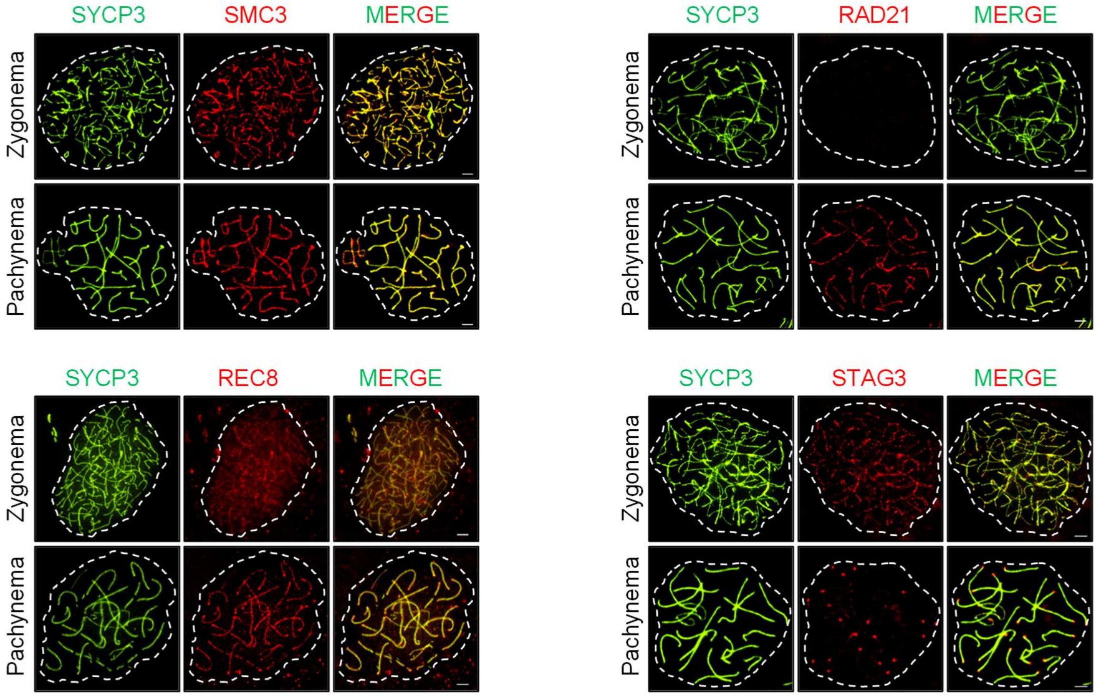


**Supplementary Figure 1. Images of cohesin components at mid-prophase I in murine meiocytes.** Images of mouse spermatocytes stained with anti-SYCP3, anti-SMC3, anti-RAD21, anti-REC8, and anti-STAG3. The distribution of cohesin components on chromosomes were observed using fluorescence microscope. Scale bars are 2.5 μm.

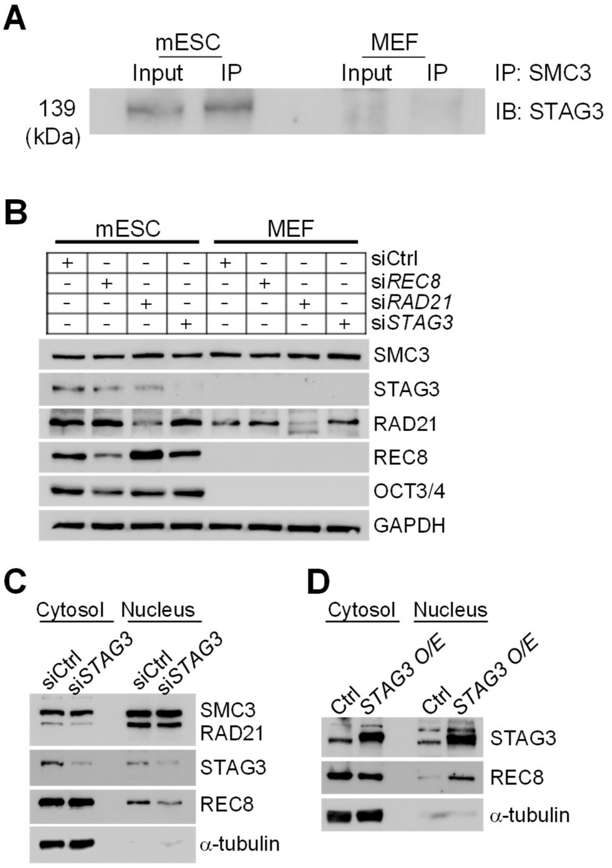


**Supplementary Figure 2. Expression and interaction of mitotic/meiotic cohesin components. (A)** Immunoprecipitation experiment in ESCs and MEFs. We pulled down SMC3 using an anti-STAG3 antibody in whole-cell extracts from ESCs and MEFs. **(B)** Knockdown of cohesin factors in ESCs and MEFs. We analyzed the expression levels of RAD21, REC8, and STAG3 following transfection with a siRNA pool against RAD21, REC8, and STAG3 (si*RAD21*, si*REC8,* and si*STAG3*). We also transfected the control cells with a nontargeting siRNA (siCtrl). GAPDH was used as a loading control. **(C and D)** ESC lysates were fractionated into cytosolic and nuclear proteins and performed immunoblot analysis. α-tubulin was used as a control for the cytosolic fraction.

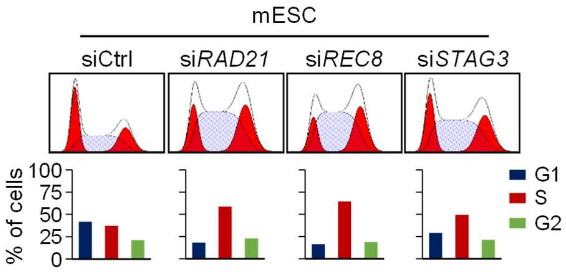


**Supplementary Figure 3. Cell cycle analysis after knockdown of cohesin components.** Cell cycle profiles of ESCs treated with si*RAD21*, si*REC8,* and si*STAG3*. We stained the cell samples with DAPI and characterized them by FACS analysis.

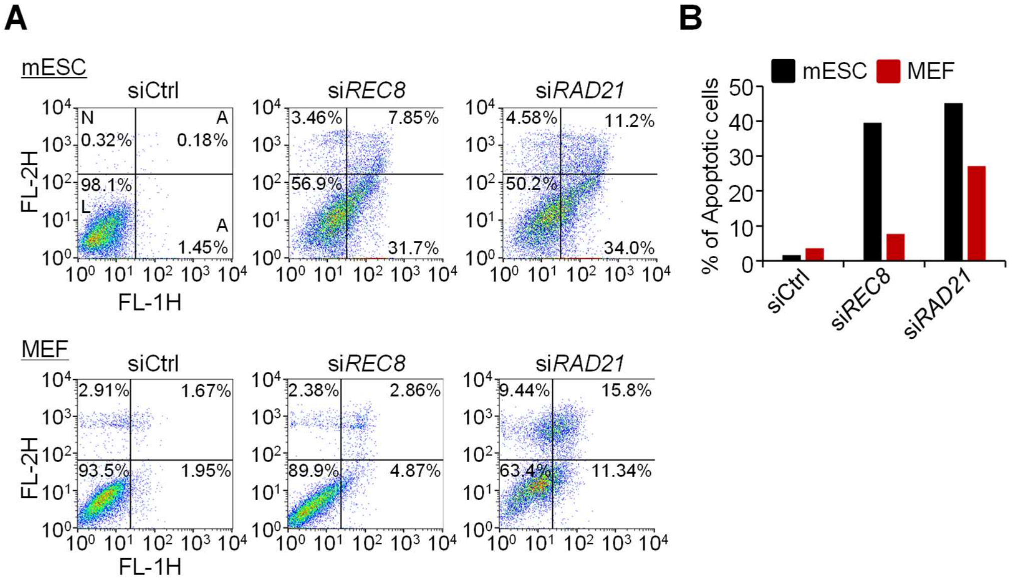


**Supplementary Figure 4. Analysis of apoptosis after knockdown of RAD21 and REC8. (A)** Analysis of apoptosis in mESCs and MEF. The cells were incubated with siRNA in serum-free medium. The proportion of apoptotic cells was quantified with FITC-conjugated annexin V (2 μg/ml) and PI (15 μg/ml). Scatter plots indicate the distribution of FITC-conjugated annexin V and PI staining. The cells are classified as "live-cell" (bottom left), "early apoptotic-cell" (bottom right), "late apoptotic-cell" (top right) and "necrotic-cell" (top left). **(B)** Quantification of apoptotic cells in mESCs and MEF.

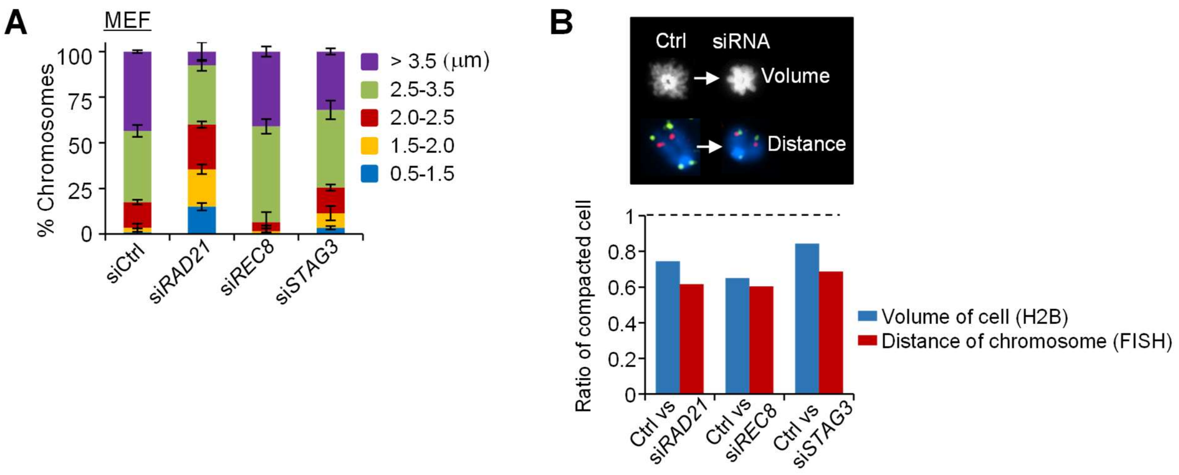


**Supplementary Figure 5. Chromosome compaction in spread samples and fixed whole nuclei during metaphase. (A)** Quantification of chromosome lengths. The length of the chromosomes in MEF was measured by calculating the distance between both sides of the telomere probes. Error bars represent mean ± SD values (N ≥ 200). **(B)** Quantification of chromosome volume in the fixed whole nuclei (H2B) and distance in spread samples hybridized with telomeric probes and locus-specific probes. The volume and distance from metaphase chromosomes in cells depleted of mitotic/meiotic cohesin components showed similar reductions in proportion.

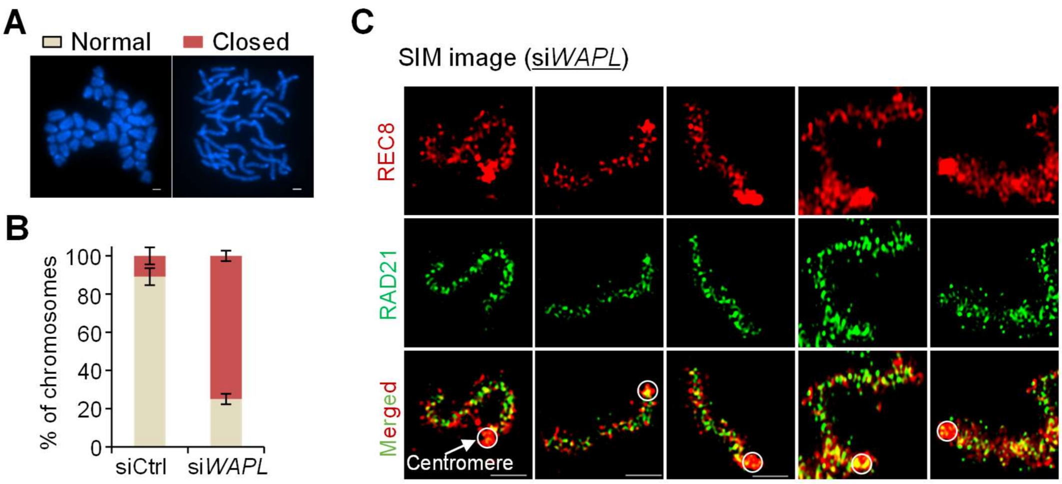


**Supplementary Figure 6. Change in cohesin localization and chromosome structure after *WAPL* knockdown. (A)** Representative images showing normal and closed chromosomes in ESCs. **(B)** Chromosome spreads of ESCs, including normal and closed chromosomes. Error bars are mean ± SD (N = 500). **(C)** Representative 3D-SIM images of WAPL-depleted cells in metaphase. Scale bars are 2.5 μm.

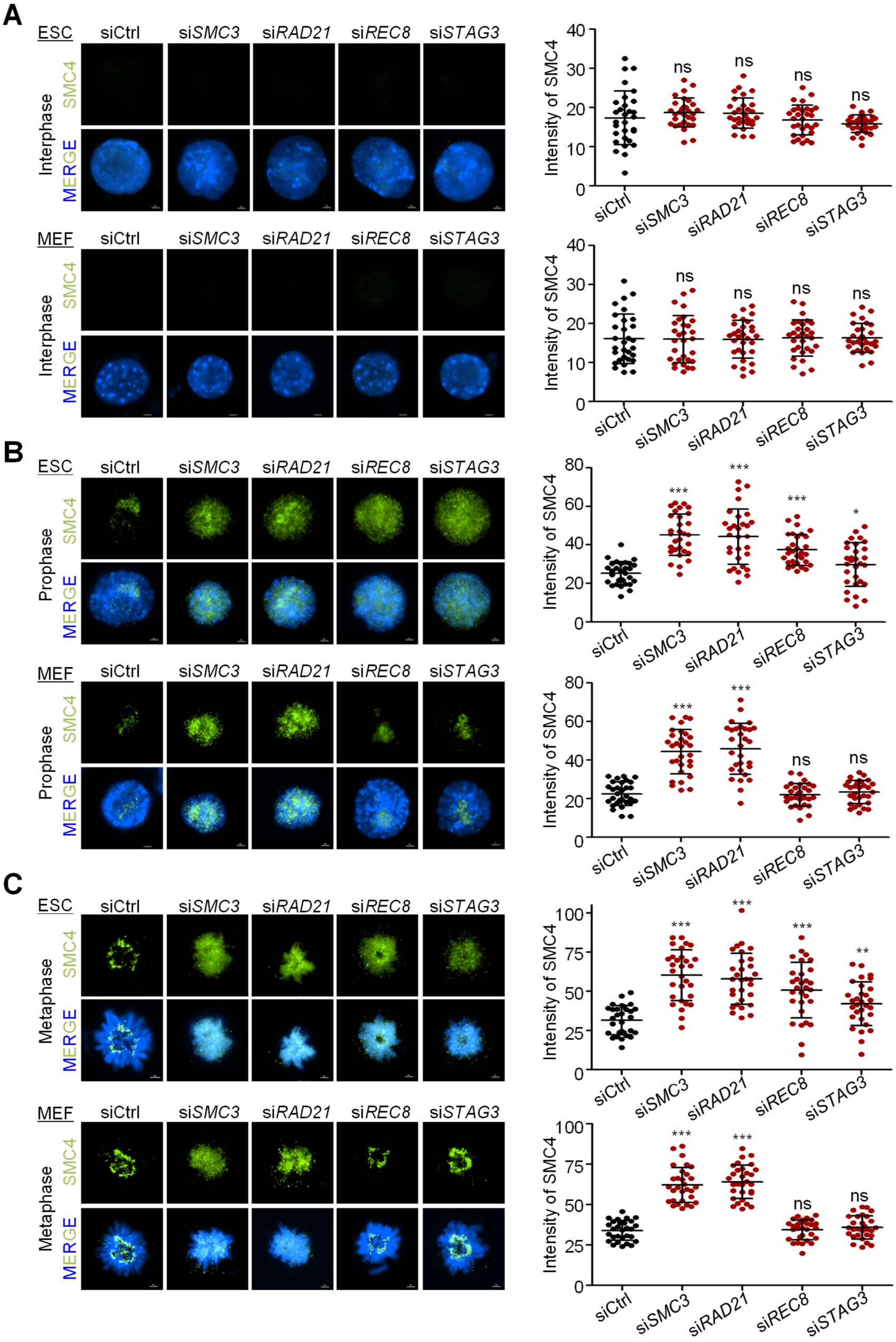

A
ESC
siCtrl
siSMC3
siRAD21
siREC8
siSTAG3
Interphase
SMC4
MERGE
Intensity of SMC4
ns
MEF
B
Prophase
***
*
C
Metaphase
**

**Supplementary Figure 7. Localization pattern of condensin from interphase to metaphase in ESCs and MEF. (A-C)** Analysis of condensin intensity from interphase to metaphase in ESCs and MEF. We immunostained interphase, prophase, and metaphase cells with an antibody against SMC4 and counterstained them with DAPI. Scale bars are 2.5 μm. We measured intensity using the Nikon NIS software. P-values (Student's t-test) were calculated using GraphPad Prism 5 software. ns: not-significant, *P < 0.5, **P < 0.01, and ***P < 0.001.

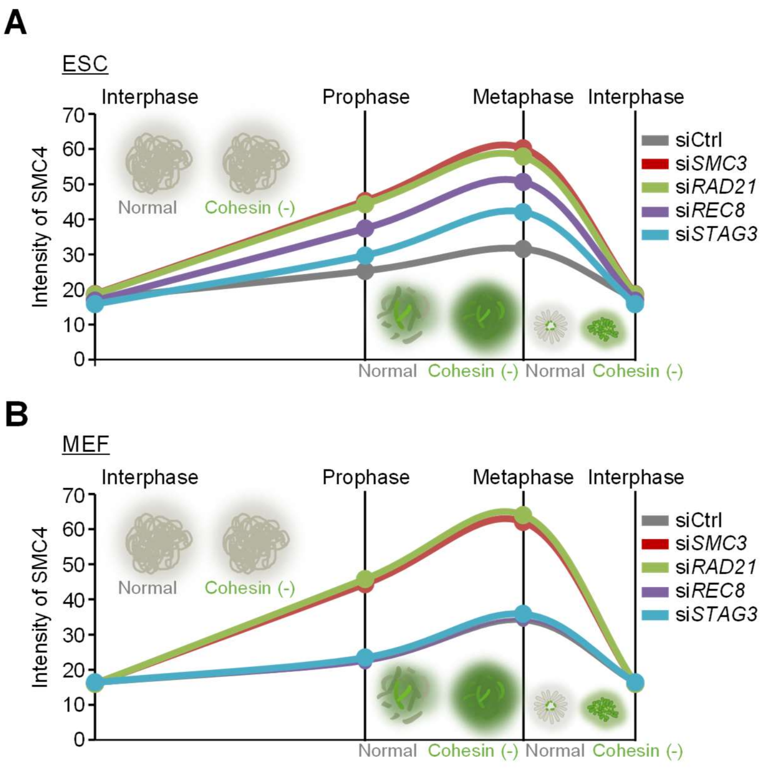


**Supplementary Figure 8. Changes in condensin intensity during the cell cycle of ESC and MEF. (A and B)** Analysis of condensin intensity at interphase, prophase, and metaphase in ESC and MEF. Intensity values of condensin were quantified from supplementary figure 7.

Supplementary Table 1. Up- or down-retulated gene list after knockdown of cohesin components.

A. Upregulated gene list after knockdown of SMC3, RAD21, and REC8.

| Gene symbol | Fold change | | | Average of normalized data (log2) | | | | Average of raw data | | | |
|---|---|---|---|---|---|---|---|---|---|---|---|
| | siRec8 /siControl | siRad21 /siControl | siSMC3 /siControl | siControl | siRec8 | siRad21 | siSMC3 | siControl | siRec8 | siRad21 | siSMC3 |
| Rb1 | 2.702 | 3.349 | 2.759 | 1.504 | 2.938 | 3.248 | 2.968 | 1.832 | 6.644 | 8.523 | 6.802 |
| Cyr61 | 6.957 | 6.534 | 7.080 | 1.343 | 4.142 | 4.051 | 4.167 | 1.533 | 16.609 | 15.629 | 16.905 |
| Sdc4 | 5.813 | 4.674 | 4.016 | 2.964 | 5.504 | 5.189 | 4.970 | 6.798 | 44.259 | 35.604 | 30.235 |
| Serpine2 | 5.304 | 6.725 | 5.395 | 1.919 | 4.326 | 4.669 | 4.351 | 2.778 | 19.013 | 24.511 | 19.341 |
| Plk2 | 5.129 | 4.518 | 3.732 | 2.774 | 5.133 | 4.950 | 4.675 | 5.831 | 34.004 | 30.006 | 24.452 |
| Mt2 | 3.233 | 2.839 | 3.785 | 7.986 | 9.679 | 9.492 | 9.907 | 252.392 | 816.647 | 721.423 | 955.671 |
| Perp | 3.138 | 3.170 | 2.889 | 3.321 | 4.970 | 4.985 | 4.851 | 8.980 | 30.271 | 30.774 | 27.770 |
| Trp53inp1 | 3.131 | 3.374 | 2.720 | 4.691 | 6.337 | 6.445 | 6.135 | 24.801 | 79.657 | 86.424 | 69.017 |
| Btg2 | 3.086 | 3.264 | 2.897 | 3.173 | 4.799 | 4.879 | 4.707 | 8.011 | 26.763 | 28.528 | 25.038 |
| Ptpn14 | 2.951 | 3.309 | 2.829 | 2.619 | 4.180 | 4.345 | 4.119 | 5.135 | 17.077 | 19.380 | 16.318 |
| Ccng1 | 2.633 | 1.817 | 1.678 | 5.605 | 7.001 | 6.466 | 6.351 | 47.605 | 126.776 | 87.722 | 80.349 |
| Gpx3 | 2.442 | 2.209 | 1.896 | 3.763 | 5.051 | 4.906 | 4.685 | 12.557 | 32.065 | 29.077 | 24.644 |
| Cald1 | 2.410 | 1.784 | 1.779 | 3.664 | 4.933 | 4.499 | 4.496 | 11.669 | 29.483 | 21.689 | 21.485 |
| Gsto1 | 2.329 | 1.626 | 1.728 | 4.308 | 5.527 | 5.009 | 5.097 | 18.793 | 45.010 | 31.306 | 33.111 |
| Mdm2 | 2.265 | 2.152 | 1.827 | 5.078 | 6.258 | 6.184 | 5.948 | 32.753 | 75.316 | 71.945 | 60.520 |
| Plod2 | 2.236 | 2.625 | 2.632 | 2.854 | 4.015 | 4.247 | 4.250 | 6.224 | 15.129 | 18.041 | 17.973 |
| Pdrg1 | 2.235 | 2.174 | 2.035 | 3.335 | 4.496 | 4.455 | 4.360 | 9.084 | 21.508 | 21.008 | 19.470 |
| Tex19.1 | 2.215 | 2.415 | 2.174 | 3.308 | 4.455 | 4.580 | 4.428 | 8.894 | 20.875 | 22.986 | 20.456 |
| Pisd-ps1 | 2.214 | 2.935 | 2.366 | 3.207 | 4.354 | 4.761 | 4.450 | 8.228 | 19.397 | 26.193 | 20.783 |
| Snhg12 | 2.186 | 2.706 | 2.521 | 3.463 | 4.591 | 4.899 | 4.797 | 10.011 | 23.039 | 28.925 | 26.701 |
| Mrgpra6 | 2.163 | 2.781 | 2.222 | 6.593 | 7.706 | 8.069 | 7.745 | 95.469 | 207.292 | 268.417 | 212.821 |
| B2m | 2.109 | 2.058 | 1.922 | 3.421 | 4.497 | 4.462 | 4.363 | 9.695 | 21.526 | 21.105 | 19.511 |
| Ptk7 | 2.056 | 2.566 | 2.298 | 4.500 | 5.540 | 5.860 | 5.701 | 21.610 | 45.418 | 57.252 | 50.840 |
| Gm11974 | 1.974 | 2.433 | 2.795 | 3.489 | 4.470 | 4.772 | 4.972 | 10.216 | 21.111 | 26.412 | 30.285 |
| Olfr856-ps1 | 1.948 | 2.805 | 1.846 | 3.809 | 4.771 | 5.297 | 4.694 | 13.005 | 26.231 | 38.442 | 24.790 |
| Rev1 | 1.947 | 1.952 | 1.757 | 3.205 | 4.166 | 4.170 | 4.018 | 8.209 | 16.903 | 17.056 | 15.150 |
| Ptp4a3 | 1.939 | 1.991 | 1.831 | 3.362 | 4.317 | 4.355 | 4.235 | 9.272 | 18.888 | 19.532 | 17.768 |
| Crip2 | 1.907 | 1.769 | 1.608 | 4.148 | 5.080 | 4.971 | 4.834 | 16.722 | 32.733 | 30.464 | 27.423 |
| Slc26a2 | 1.869 | 1.812 | 2.017 | 3.185 | 4.087 | 4.042 | 4.197 | 8.085 | 15.958 | 15.530 | 17.277 |
| Susd6 | 1.867 | 2.003 | 1.867 | 3.249 | 4.150 | 4.252 | 4.150 | 8.500 | 16.713 | 18.107 | 16.701 |
| S100a11 | 1.863 | 1.816 | 2.024 | 5.827 | 6.725 | 6.688 | 6.845 | 55.730 | 104.529 | 102.454 | 113.519 |
| Ece1 | 1.858 | 2.138 | 1.819 | 3.727 | 4.621 | 4.823 | 4.590 | 12.228 | 23.541 | 27.397 | 23.012 |
| Lama5 | 1.841 | 3.223 | 2.630 | 3.219 | 4.099 | 4.907 | 4.614 | 8.300 | 16.095 | 29.098 | 23.403 |
| Ccp110 | 1.813 | 1.821 | 1.948 | 3.507 | 4.365 | 4.372 | 4.469 | 10.358 | 19.558 | 19.765 | 21.075 |
| Rbms1 | 1.783 | 1.817 | 1.983 | 3.205 | 4.040 | 4.067 | 4.193 | 8.217 | 15.406 | 15.808 | 17.227 |
| Arl6ip5 | 1.727 | 1.720 | 2.002 | 4.034 | 4.822 | 4.816 | 5.035 | 15.367 | 27.205 | 27.266 | 31.674 |
| Tnpo2 | 1.721 | 2.065 | 1.940 | 4.037 | 4.820 | 5.084 | 4.993 | 15.405 | 27.180 | 33.016 | 30.747 |
| Eda2r | 1.709 | 2.223 | 2.165 | 3.281 | 4.054 | 4.434 | 4.396 | 8.713 | 15.569 | 20.685 | 19.988 |
| Mbnl2 | 1.695 | 2.117 | 1.878 | 3.627 | 4.389 | 4.709 | 4.536 | 11.346 | 19.893 | 25.241 | 22.132 |
| Paip2b | 1.673 | 1.887 | 1.848 | 4.356 | 5.098 | 5.272 | 5.242 | 19.455 | 33.167 | 37.766 | 36.726 |
| Serpinh1 | 1.653 | 2.570 | 2.221 | 4.057 | 4.781 | 5.418 | 5.208 | 15.624 | 26.426 | 41.895 | 35.842 |
| Ptgr1 | 1.648 | 1.646 | 1.618 | 3.766 | 4.487 | 4.486 | 4.461 | 12.594 | 21.376 | 21.475 | 20.950 |
| Ifrd1 | 1.628 | 2.138 | 2.076 | 4.392 | 5.095 | 5.488 | 5.445 | 19.973 | 33.085 | 44.020 | 42.423 |
| Plin2 | 1.589 | 1.712 | 1.521 | 3.448 | 4.117 | 4.224 | 4.053 | 9.909 | 16.308 | 17.747 | 15.547 |
| Tmtc3 | 1.578 | 1.645 | 1.575 | 3.608 | 4.266 | 4.327 | 4.263 | 11.181 | 18.192 | 19.129 | 18.136 |
| Cldn6 | 1.575 | 1.595 | 1.548 | 3.774 | 4.430 | 4.448 | 4.404 | 12.665 | 20.496 | 20.892 | 20.105 |
| Anxa2 | 1.567 | 2.773 | 2.635 | 3.852 | 4.500 | 5.323 | 5.250 | 13.418 | 21.560 | 39.152 | 36.914 |
| Stk25 | 1.553 | 2.230 | 1.989 | 3.392 | 4.027 | 4.549 | 4.384 | 9.486 | 15.261 | 22.480 | 19.818 |
| Hspg2 | 1.522 | 3.283 | 2.255 | 3.905 | 4.511 | 5.620 | 5.078 | 13.964 | 21.734 | 48.328 | 32.662 |
| Laptm4a | 1.517 | 1.514 | 1.552 | 5.207 | 5.808 | 5.805 | 5.841 | 35.898 | 54.864 | 55.094 | 56.126 |

B. Downregulated gene list after knockdown of SMC3, RAD21, and REC8.

| Gene symbol | Fold change | | | Average of normalized data (log2) | | | | Average of raw data | | | |
|---|---|---|---|---|---|---|---|---|---|---|---|
| | siRec8 /siControl | siRad21 /siControl | siSMC3 /siControl | siControl | siRec8 | siRad21 | siSMC3 | siControl | siRec8 | siRad21 | siSMC3 |
| Oaz1-ps | 0.474 | 0.004 | 0.047 | 7.818 | 6.741 | 0.006 | 3.420 | 224.432 | 105.700 | 0.009 | 9.648 |
| Med8 | 0.666 | 0.546 | 0.630 | 5.306 | 4.720 | 4.432 | 4.639 | 38.519 | 25.289 | 20.663 | 23.822 |
| Slc25a5 | 0.666 | 0.405 | 0.515 | 9.367 | 8.781 | 8.063 | 8.410 | 658.866 | 437.633 | 267.430 | 337.859 |
| Rps14 | 0.665 | 0.445 | 0.510 | 12.563 | 11.976 | 11.395 | 11.591 | 6046.390 | 4015.820 | 2700.600 | 3073.470 |
| Nr2f6 | 0.665 | 0.644 | 0.659 | 4.722 | 4.134 | 4.087 | 4.121 | 25.375 | 16.513 | 16.045 | 16.333 |
| Eif2d | 0.665 | 0.502 | 0.588 | 7.808 | 7.220 | 6.813 | 7.042 | 222.933 | 147.680 | 111.859 | 130.254 |
| Ndufaf3 | 0.664 | 0.646 | 0.647 | 4.369 | 3.779 | 3.740 | 3.741 | 19.652 | 12.695 | 12.403 | 12.325 |
| Ppa1 | 0.664 | 0.468 | 0.533 | 7.999 | 7.409 | 6.905 | 7.093 | 254.664 | 168.462 | 119.261 | 134.962 |
| AI837181 | 0.664 | 0.478 | 0.606 | 4.974 | 4.384 | 3.909 | 4.251 | 30.408 | 19.819 | 14.072 | 17.974 |
| Msto1 | 0.664 | 0.383 | 0.500 | 6.223 | 5.631 | 4.837 | 5.222 | 73.623 | 48.443 | 27.691 | 36.168 |
| Cct6a | 0.663 | 0.455 | 0.526 | 9.349 | 8.757 | 8.215 | 8.424 | 650.784 | 430.614 | 297.136 | 341.132 |
| Mbd3 | 0.663 | 0.563 | 0.567 | 6.818 | 6.226 | 5.990 | 6.001 | 111.785 | 73.645 | 62.772 | 62.787 |
| Eif2s2 | 0.663 | 0.424 | 0.531 | 8.270 | 7.677 | 7.034 | 7.356 | 307.461 | 203.126 | 130.477 | 162.178 |
| Cuedc2 | 0.662 | 0.588 | 0.647 | 6.121 | 5.527 | 5.356 | 5.494 | 68.550 | 44.987 | 40.095 | 43.898 |
| Ndufs7 | 0.662 | 0.599 | 0.611 | 6.972 | 6.377 | 6.233 | 6.261 | 124.404 | 81.880 | 74.468 | 75.417 |
| Aven | 0.662 | 0.539 | 0.638 | 5.747 | 5.152 | 4.854 | 5.098 | 52.659 | 34.464 | 28.022 | 33.113 |
| Tkt | 0.662 | 0.595 | 0.641 | 8.097 | 7.501 | 7.348 | 7.454 | 272.466 | 179.641 | 162.467 | 173.677 |
| Rassf7 | 0.662 | 0.462 | 0.575 | 4.688 | 4.092 | 3.574 | 3.889 | 24.756 | 16.010 | 10.951 | 13.762 |
| Bysl | 0.660 | 0.437 | 0.539 | 5.884 | 5.285 | 4.691 | 4.991 | 58.004 | 37.890 | 24.925 | 30.686 |
| Smpd2 | 0.660 | 0.460 | 0.547 | 4.270 | 3.671 | 3.149 | 3.401 | 18.281 | 11.705 | 7.903 | 9.524 |
| Tead2 | 0.660 | 0.573 | 0.619 | 6.358 | 5.758 | 5.555 | 5.666 | 80.961 | 52.969 | 46.166 | 49.583 |
| Maz | 0.659 | 0.559 | 0.646 | 7.329 | 6.728 | 6.489 | 6.699 | 159.641 | 104.707 | 89.124 | 102.540 |
| Rps3 | 0.658 | 0.529 | 0.584 | 10.668 | 10.065 | 9.749 | 9.893 | 1624.680 | 1067.465 | 862.603 | 946.001 |
| Ndufs8 | 0.658 | 0.606 | 0.638 | 6.952 | 6.349 | 6.229 | 6.304 | 122.704 | 80.285 | 74.276 | 77.741 |
| Pagr1a | 0.658 | 0.507 | 0.614 | 5.553 | 4.949 | 4.572 | 4.849 | 45.910 | 29.798 | 22.863 | 27.721 |
| Basp1 | 0.657 | 0.566 | 0.627 | 7.237 | 6.632 | 6.417 | 6.565 | 149.742 | 97.893 | 84.724 | 93.309 |
| Atic | 0.657 | 0.512 | 0.606 | 7.834 | 7.228 | 6.867 | 7.110 | 226.945 | 148.488 | 116.119 | 136.653 |
| Ssbp4 | 0.657 | 0.568 | 0.641 | 6.707 | 6.101 | 5.892 | 6.065 | 103.392 | 67.448 | 58.571 | 65.729 |
| Tfpt | 0.657 | 0.419 | 0.506 | 5.207 | 4.600 | 3.953 | 4.224 | 35.891 | 23.188 | 14.536 | 17.618 |
| Taf10 | 0.656 | 0.647 | 0.656 | 7.341 | 6.734 | 6.714 | 6.734 | 161.021 | 105.133 | 104.348 | 105.030 |
| Pold2 | 0.656 | 0.516 | 0.583 | 7.003 | 6.395 | 6.049 | 6.225 | 127.160 | 82.955 | 65.428 | 73.504 |
| Grtp1 | 0.656 | 0.640 | 0.651 | 6.028 | 5.419 | 5.383 | 5.409 | 64.188 | 41.669 | 40.865 | 41.354 |
| Psph | 0.655 | 0.587 | 0.604 | 5.672 | 5.062 | 4.905 | 4.944 | 49.945 | 32.310 | 29.056 | 29.679 |
| Trmt112 | 0.655 | 0.497 | 0.535 | 7.271 | 6.660 | 6.262 | 6.369 | 153.324 | 99.842 | 76.013 | 81.345 |
| Pih1d1 | 0.655 | 0.452 | 0.497 | 6.187 | 5.576 | 5.041 | 5.178 | 71.798 | 46.573 | 32.038 | 35.053 |
| Mrpl11 | 0.654 | 0.428 | 0.508 | 5.403 | 4.790 | 4.179 | 4.426 | 41.266 | 26.590 | 17.182 | 20.421 |
| Smarcb1 | 0.654 | 0.482 | 0.575 | 6.982 | 6.369 | 5.929 | 6.183 | 125.273 | 81.413 | 60.140 | 71.378 |
| Stub1 | 0.653 | 0.542 | 0.620 | 6.512 | 5.898 | 5.629 | 5.822 | 90.149 | 58.463 | 48.643 | 55.384 |
| Arl6ip4 | 0.653 | 0.440 | 0.550 | 6.098 | 5.484 | 4.913 | 5.237 | 67.453 | 43.628 | 29.232 | 36.561 |
| Timm23 | 0.653 | 0.442 | 0.558 | 8.228 | 7.612 | 7.048 | 7.387 | 298.490 | 194.158 | 131.821 | 165.695 |
| Psmd13 | 0.653 | 0.566 | 0.657 | 7.544 | 6.928 | 6.722 | 6.939 | 185.468 | 120.471 | 104.945 | 121.216 |
| Eif3d | 0.652 | 0.475 | 0.569 | 8.208 | 7.591 | 7.134 | 7.394 | 294.338 | 191.321 | 139.924 | 166.544 |
| Rab8a | 0.652 | 0.523 | 0.651 | 5.790 | 5.173 | 4.855 | 5.171 | 54.283 | 34.986 | 28.038 | 34.897 |
| Sra1 | 0.651 | 0.563 | 0.647 | 6.408 | 5.787 | 5.579 | 5.779 | 83.830 | 54.082 | 46.960 | 53.699 |
| Hnrnpab | 0.650 | 0.425 | 0.554 | 9.477 | 8.857 | 8.244 | 8.626 | 711.120 | 461.300 | 303.145 | 392.603 |
| Rnaseh2a | 0.650 | 0.517 | 0.579 | 6.014 | 5.391 | 5.062 | 5.225 | 63.544 | 40.855 | 32.517 | 36.259 |
| Rpl30 | 0.648 | 0.368 | 0.441 | 8.298 | 7.672 | 6.856 | 7.116 | 313.455 | 202.459 | 115.238 | 137.159 |
| Tsen34 | 0.648 | 0.508 | 0.599 | 7.073 | 6.446 | 6.097 | 6.333 | 133.479 | 85.955 | 67.669 | 79.331 |
| Dohh | 0.647 | 0.464 | 0.529 | 6.807 | 6.180 | 5.698 | 5.888 | 110.895 | 71.305 | 51.086 | 57.990 |
| Ubxn6 | 0.646 | 0.558 | 0.571 | 5.326 | 4.696 | 4.484 | 4.517 | 39.081 | 24.854 | 21.458 | 21.817 |
| Gltscr2 | 0.646 | 0.397 | 0.490 | 8.156 | 7.526 | 6.823 | 7.127 | 283.887 | 182.778 | 112.647 | 138.219 |
| Anp32b | 0.646 | 0.535 | 0.578 | 9.770 | 9.140 | 8.867 | 8.980 | 871.179 | 561.650 | 467.548 | 502.084 |
| Sphk2 | 0.646 | 0.485 | 0.605 | 4.090 | 3.459 | 3.046 | 3.365 | 16.018 | 9.973 | 7.287 | 9.264 |
| Isyna1 | 0.646 | 0.523 | 0.617 | 6.895 | 6.264 | 5.960 | 6.199 | 117.927 | 75.662 | 61.447 | 72.200 |

| | | | | | | | | | | | |
|---|---|---|---|---|---|---|---|---|---|---|---|
| Chchd1 | 0.645 | 0.579 | 0.627 | 7.248 | 6.615 | 6.460 | 6.575 | 150.824 | 96.728 | 87.307 | 93.967 |
| Ruvbl1 | 0.645 | 0.451 | 0.549 | 7.722 | 7.088 | 6.572 | 6.858 | 209.943 | 134.712 | 94.479 | 114.529 |
| Rabif | 0.644 | 0.559 | 0.636 | 5.890 | 5.254 | 5.051 | 5.236 | 58.238 | 37.063 | 32.274 | 36.542 |
| Nme6 | 0.644 | 0.491 | 0.539 | 5.109 | 4.473 | 4.084 | 4.219 | 33.488 | 21.155 | 16.017 | 17.552 |
| Rnf187 | 0.643 | 0.423 | 0.506 | 7.455 | 6.819 | 6.213 | 6.472 | 174.339 | 111.583 | 73.438 | 87.419 |
| Sgk1 | 0.642 | 0.427 | 0.648 | 8.199 | 7.560 | 6.969 | 7.573 | 292.553 | 187.200 | 124.752 | 188.702 |
| Yif1b | 0.641 | 0.583 | 0.666 | 6.320 | 5.678 | 5.541 | 5.734 | 78.808 | 50.069 | 45.713 | 52.020 |
| Rdh11 | 0.640 | 0.657 | 0.658 | 4.634 | 3.991 | 4.029 | 4.031 | 23.817 | 14.859 | 15.374 | 15.289 |
| Gm4737 | 0.640 | 0.468 | 0.586 | 6.017 | 5.373 | 4.921 | 5.245 | 63.691 | 40.323 | 29.396 | 36.788 |
| Eif2b4 | 0.639 | 0.376 | 0.522 | 6.152 | 5.505 | 4.741 | 5.213 | 70.055 | 44.290 | 25.837 | 35.948 |
| Ube2r2 | 0.638 | 0.627 | 0.590 | 5.556 | 4.907 | 4.882 | 4.795 | 46.013 | 28.930 | 28.584 | 26.669 |
| Tbcb | 0.638 | 0.329 | 0.481 | 6.806 | 6.156 | 5.204 | 5.751 | 110.770 | 70.144 | 35.989 | 52.652 |
| Serbp1 | 0.636 | 0.520 | 0.604 | 8.334 | 7.682 | 7.391 | 7.606 | 321.302 | 203.744 | 167.399 | 193.034 |
| Csrp2 | 0.635 | 0.401 | 0.499 | 8.353 | 7.699 | 7.034 | 7.350 | 325.698 | 206.179 | 130.503 | 161.554 |
| Nr0b1 | 0.635 | 0.406 | 0.561 | 7.002 | 6.347 | 5.702 | 6.168 | 127.093 | 80.199 | 51.224 | 70.625 |
| Apex1 | 0.634 | 0.352 | 0.500 | 6.629 | 5.972 | 5.122 | 5.631 | 97.913 | 61.610 | 33.949 | 48.359 |
| Pmvk | 0.633 | 0.563 | 0.570 | 6.384 | 5.724 | 5.555 | 5.573 | 82.424 | 51.732 | 46.183 | 46.424 |
| Gjb5 | 0.633 | 0.349 | 0.384 | 4.777 | 4.116 | 3.258 | 3.397 | 26.385 | 16.303 | 8.602 | 9.497 |
| Tgif1 | 0.633 | 0.480 | 0.629 | 7.377 | 6.717 | 6.319 | 6.709 | 165.106 | 103.921 | 79.099 | 103.251 |
| Carkd | 0.633 | 0.522 | 0.655 | 4.350 | 3.690 | 3.413 | 3.739 | 19.383 | 11.872 | 9.686 | 12.309 |
| Uqcc2 | 0.631 | 0.597 | 0.633 | 8.393 | 7.729 | 7.649 | 7.734 | 334.899 | 210.581 | 200.454 | 211.123 |
| Srm | 0.631 | 0.363 | 0.446 | 8.185 | 7.520 | 6.724 | 7.020 | 289.673 | 182.018 | 105.097 | 128.306 |
| Bckdk | 0.631 | 0.523 | 0.612 | 5.154 | 4.489 | 4.218 | 4.444 | 34.572 | 21.390 | 17.674 | 20.692 |
| Rangrf | 0.630 | 0.382 | 0.461 | 7.154 | 6.487 | 5.764 | 6.038 | 141.308 | 88.477 | 53.542 | 64.460 |
| Ict1 | 0.630 | 0.403 | 0.464 | 6.493 | 5.826 | 5.183 | 5.385 | 89.022 | 55.570 | 35.464 | 40.621 |
| Nubp1 | 0.629 | 0.471 | 0.599 | 5.560 | 4.891 | 4.475 | 4.820 | 46.135 | 28.594 | 21.316 | 27.140 |
| Psmb5 | 0.629 | 0.515 | 0.579 | 8.701 | 8.032 | 7.744 | 7.912 | 414.815 | 260.010 | 214.188 | 238.876 |
| Ppif | 0.627 | 0.473 | 0.548 | 6.287 | 5.613 | 5.208 | 5.420 | 77.018 | 47.811 | 36.085 | 41.645 |
| Lap3 | 0.627 | 0.406 | 0.485 | 8.222 | 7.548 | 6.920 | 7.179 | 297.272 | 185.583 | 120.503 | 143.329 |
| Ahcy | 0.626 | 0.439 | 0.557 | 6.648 | 5.973 | 5.461 | 5.804 | 99.222 | 61.655 | 43.212 | 54.674 |
| Smarcd3 | 0.626 | 0.559 | 0.510 | 5.324 | 4.648 | 4.485 | 4.353 | 39.036 | 24.007 | 21.462 | 19.356 |
| H2afz | 0.625 | 0.411 | 0.533 | 10.534 | 9.855 | 9.251 | 9.626 | 1480.770 | 922.698 | 610.214 | 786.346 |
| Mars | 0.624 | 0.620 | 0.662 | 6.856 | 6.176 | 6.168 | 6.262 | 114.770 | 71.089 | 71.143 | 75.447 |
| Tmem160 | 0.623 | 0.441 | 0.461 | 6.419 | 5.738 | 5.239 | 5.302 | 84.532 | 52.223 | 36.904 | 38.309 |
| Ube2m | 0.623 | 0.372 | 0.494 | 7.757 | 7.075 | 6.329 | 6.740 | 215.187 | 133.472 | 79.693 | 105.474 |
| Mrto4 | 0.623 | 0.496 | 0.560 | 7.220 | 6.538 | 6.207 | 6.383 | 147.982 | 91.659 | 73.150 | 82.126 |
| Gadd45gip1 | 0.622 | 0.465 | 0.509 | 4.963 | 4.279 | 3.857 | 3.987 | 30.154 | 18.361 | 13.542 | 14.802 |
| Fiz1 | 0.621 | 0.412 | 0.532 | 5.983 | 5.296 | 4.703 | 5.073 | 62.194 | 38.192 | 25.131 | 32.538 |
| Samm50 | 0.621 | 0.481 | 0.568 | 8.275 | 7.588 | 7.219 | 7.460 | 308.476 | 190.947 | 148.481 | 174.397 |
| Mpnd | 0.621 | 0.594 | 0.651 | 5.899 | 5.211 | 5.147 | 5.278 | 58.619 | 35.938 | 34.549 | 37.672 |
| Tfeb | 0.621 | 0.407 | 0.558 | 5.441 | 4.753 | 4.145 | 4.598 | 42.392 | 25.888 | 16.751 | 23.126 |
| Cops6 | 0.620 | 0.433 | 0.464 | 8.454 | 7.765 | 7.245 | 7.345 | 349.283 | 215.892 | 151.182 | 160.969 |
| Rpl7a | 0.620 | 0.475 | 0.573 | 11.805 | 11.116 | 10.731 | 11.001 | 3574.160 | 2211.820 | 1704.995 | 2040.590 |
| Pgls | 0.620 | 0.630 | 0.557 | 6.911 | 6.221 | 6.244 | 6.066 | 119.257 | 73.375 | 75.052 | 65.773 |
| Acat2 | 0.619 | 0.490 | 0.499 | 6.495 | 5.803 | 5.464 | 5.492 | 89.092 | 54.678 | 43.304 | 43.827 |
| Gpa33 | 0.619 | 0.521 | 0.521 | 5.533 | 4.841 | 4.592 | 4.594 | 45.275 | 27.586 | 23.208 | 23.063 |
| Dkc1 | 0.617 | 0.487 | 0.551 | 7.685 | 6.989 | 6.647 | 6.826 | 204.649 | 125.719 | 99.551 | 112.001 |
| Rpl18a | 0.617 | 0.500 | 0.551 | 10.421 | 9.724 | 9.422 | 9.561 | 1368.930 | 842.674 | 687.363 | 751.442 |
| Prmt6 | 0.616 | 0.486 | 0.618 | 4.018 | 3.319 | 2.979 | 3.325 | 15.193 | 8.957 | 6.908 | 8.985 |
| Uxt | 0.615 | 0.605 | 0.641 | 5.023 | 4.322 | 4.297 | 4.380 | 31.478 | 18.953 | 18.721 | 19.752 |
| Hist2h2bb | 0.615 | 0.448 | 0.596 | 6.497 | 5.796 | 5.338 | 5.752 | 89.271 | 54.433 | 39.588 | 52.696 |
| Tubb3 | 0.615 | 0.551 | 0.545 | 4.096 | 3.395 | 3.236 | 3.221 | 16.092 | 9.492 | 8.454 | 8.290 |
| Dph3 | 0.615 | 0.508 | 0.590 | 4.381 | 3.679 | 3.404 | 3.620 | 19.810 | 11.776 | 9.618 | 11.252 |
| Tmem11 | 0.615 | 0.486 | 0.592 | 6.207 | 5.506 | 5.166 | 5.452 | 72.828 | 44.316 | 35.028 | 42.614 |
| Agpat2 | 0.615 | 0.598 | 0.614 | 5.044 | 4.342 | 4.302 | 4.340 | 31.968 | 19.230 | 18.792 | 19.177 |
| Nudt8 | 0.613 | 0.500 | 0.510 | 4.708 | 4.003 | 3.707 | 3.737 | 25.122 | 14.990 | 12.107 | 12.288 |

| | | | | | | | | | | | |
|---|---|---|---|---|---|---|---|---|---|---|---|
| Cyc1 | 0.609 | 0.505 | 0.583 | 8.375 | 7.660 | 7.390 | 7.596 | 330.646 | 200.667 | 167.307 | 191.766 |
| Tars | 0.605 | 0.550 | 0.600 | 6.918 | 6.194 | 6.056 | 6.181 | 119.827 | 71.997 | 65.752 | 71.302 |
| 2810428I15Rik | 0.605 | 0.517 | 0.612 | 6.856 | 6.132 | 5.903 | 6.148 | 114.765 | 68.934 | 59.058 | 69.634 |
| Snx17 | 0.605 | 0.532 | 0.562 | 6.014 | 5.289 | 5.103 | 5.183 | 63.560 | 37.983 | 33.490 | 35.187 |
| Phb2 | 0.605 | 0.431 | 0.493 | 9.153 | 8.428 | 7.937 | 8.133 | 567.823 | 342.390 | 244.942 | 278.677 |
| Tmem234 | 0.605 | 0.512 | 0.568 | 6.959 | 6.233 | 5.993 | 6.143 | 123.278 | 74.012 | 62.902 | 69.396 |
| Plp2 | 0.604 | 0.521 | 0.533 | 6.923 | 6.197 | 5.982 | 6.015 | 120.247 | 72.162 | 62.434 | 63.445 |
| Mvk | 0.604 | 0.489 | 0.509 | 5.320 | 4.592 | 4.287 | 4.347 | 38.918 | 23.059 | 18.585 | 19.270 |
| Tuba1b | 0.604 | 0.400 | 0.473 | 9.581 | 8.853 | 8.258 | 8.502 | 764.021 | 460.191 | 306.256 | 360.066 |
| Nsun5 | 0.604 | 0.413 | 0.510 | 4.418 | 3.689 | 3.140 | 3.445 | 20.356 | 11.867 | 7.847 | 9.849 |
| Slc25a1 | 0.602 | 0.442 | 0.517 | 6.398 | 5.666 | 5.218 | 5.447 | 83.234 | 49.623 | 36.361 | 42.452 |
| Mepce | 0.601 | 0.476 | 0.571 | 4.798 | 4.064 | 3.728 | 3.989 | 26.798 | 15.681 | 12.299 | 14.821 |
| Fdft1 | 0.600 | 0.541 | 0.524 | 5.838 | 5.101 | 4.951 | 4.904 | 56.142 | 33.236 | 30.043 | 28.834 |
| Abhd17a | 0.598 | 0.493 | 0.566 | 6.501 | 5.759 | 5.479 | 5.679 | 89.484 | 53.008 | 43.759 | 50.051 |
| Tomm40 | 0.597 | 0.403 | 0.508 | 7.951 | 7.207 | 6.638 | 6.973 | 246.229 | 146.365 | 98.968 | 124.119 |
| Tpm3 | 0.597 | 0.439 | 0.577 | 8.633 | 7.888 | 7.446 | 7.838 | 395.607 | 235.285 | 173.955 | 226.975 |
| Hax1 | 0.597 | 0.515 | 0.556 | 7.349 | 6.604 | 6.392 | 6.503 | 161.927 | 96.033 | 83.267 | 89.383 |
| Arhgdia | 0.596 | 0.477 | 0.561 | 7.777 | 7.031 | 6.709 | 6.944 | 218.143 | 129.434 | 103.982 | 121.616 |
| Thap11 | 0.594 | 0.469 | 0.584 | 5.350 | 4.600 | 4.257 | 4.575 | 39.762 | 23.188 | 18.193 | 22.746 |
| Timm13 | 0.594 | 0.427 | 0.481 | 6.782 | 6.030 | 5.556 | 5.727 | 108.945 | 64.182 | 46.200 | 51.774 |
| Naa10 | 0.592 | 0.340 | 0.451 | 7.361 | 6.604 | 5.804 | 6.211 | 163.239 | 96.029 | 55.079 | 72.798 |
| Pycr2 | 0.592 | 0.408 | 0.575 | 6.884 | 6.126 | 5.590 | 6.085 | 116.997 | 68.672 | 47.334 | 66.638 |
| Rsph1 | 0.591 | 0.387 | 0.466 | 4.401 | 3.643 | 3.031 | 3.299 | 20.106 | 11.460 | 7.201 | 8.804 |
| Mybl2 | 0.589 | 0.420 | 0.495 | 9.296 | 8.534 | 8.043 | 8.283 | 627.187 | 368.617 | 263.740 | 309.243 |
| Ccdc22 | 0.587 | 0.526 | 0.611 | 4.432 | 3.664 | 3.505 | 3.721 | 20.567 | 11.641 | 10.388 | 12.140 |
| Nelfe | 0.587 | 0.402 | 0.530 | 6.550 | 5.781 | 5.235 | 5.633 | 92.606 | 53.855 | 36.804 | 48.440 |
| Tsta3 | 0.586 | 0.417 | 0.513 | 6.041 | 5.271 | 4.778 | 5.079 | 64.805 | 37.503 | 26.536 | 32.667 |
| Ak2 | 0.586 | 0.553 | 0.625 | 7.831 | 7.060 | 6.977 | 7.152 | 226.431 | 132.068 | 125.387 | 140.717 |
| Cenpb | 0.586 | 0.421 | 0.459 | 6.127 | 5.355 | 4.879 | 5.004 | 68.815 | 39.811 | 28.521 | 30.979 |
| Tuba1a | 0.585 | 0.429 | 0.449 | 6.876 | 6.103 | 5.654 | 5.720 | 116.301 | 67.535 | 49.540 | 51.495 |
| Fam213b | 0.584 | 0.546 | 0.510 | 5.488 | 4.713 | 4.613 | 4.518 | 43.838 | 25.150 | 23.560 | 21.821 |
| Inhbb | 0.584 | 0.498 | 0.500 | 6.290 | 5.514 | 5.283 | 5.290 | 77.168 | 44.568 | 38.075 | 37.976 |
| Map2k2 | 0.583 | 0.416 | 0.526 | 6.484 | 5.707 | 5.217 | 5.557 | 88.452 | 51.094 | 36.335 | 45.902 |
| Fkbp4 | 0.583 | 0.375 | 0.437 | 9.390 | 8.611 | 7.976 | 8.196 | 669.276 | 389.026 | 251.562 | 291.218 |
| Cisd3 | 0.582 | 0.417 | 0.450 | 7.779 | 6.999 | 6.519 | 6.629 | 218.458 | 126.607 | 91.025 | 97.591 |
| Mrpl12 | 0.582 | 0.344 | 0.469 | 8.081 | 7.300 | 6.542 | 6.988 | 269.539 | 156.166 | 92.484 | 125.425 |
| Hist1h4i | 0.582 | 0.530 | 0.511 | 5.130 | 4.349 | 4.215 | 4.162 | 34.011 | 19.327 | 17.634 | 16.832 |
| Galk1 | 0.581 | 0.495 | 0.548 | 7.490 | 6.707 | 6.475 | 6.621 | 178.600 | 103.226 | 88.275 | 97.082 |
| Epb4.1l4a | 0.580 | 0.418 | 0.505 | 5.716 | 4.930 | 4.459 | 4.729 | 51.527 | 29.396 | 21.074 | 25.429 |
| Eif6 | 0.579 | 0.357 | 0.453 | 8.329 | 7.541 | 6.841 | 7.185 | 320.224 | 184.761 | 114.038 | 143.972 |
| Gdf3 | 0.577 | 0.435 | 0.607 | 5.529 | 4.736 | 4.328 | 4.809 | 45.132 | 25.586 | 19.159 | 26.937 |
| Mvb12a | 0.577 | 0.372 | 0.475 | 6.157 | 5.363 | 4.731 | 5.083 | 70.277 | 40.034 | 25.659 | 32.764 |
| Alkbh7 | 0.576 | 0.552 | 0.478 | 5.439 | 4.643 | 4.583 | 4.374 | 42.367 | 23.918 | 23.049 | 19.660 |
| Hist1h4k | 0.575 | 0.397 | 0.543 | 5.840 | 5.042 | 4.507 | 4.958 | 56.222 | 31.860 | 21.822 | 29.974 |
| Ufc1 | 0.575 | 0.507 | 0.542 | 7.206 | 6.406 | 6.227 | 6.321 | 146.517 | 83.591 | 74.143 | 78.674 |
| Qdpr | 0.570 | 0.529 | 0.539 | 7.046 | 6.236 | 6.126 | 6.155 | 130.997 | 74.174 | 69.100 | 69.981 |
| Mical1 | 0.565 | 0.529 | 0.644 | 4.262 | 3.439 | 3.342 | 3.628 | 18.175 | 9.814 | 9.177 | 11.320 |
| Ctu2 | 0.565 | 0.455 | 0.558 | 4.445 | 3.621 | 3.308 | 3.604 | 20.768 | 11.268 | 8.937 | 11.113 |
| Exosc5 | 0.562 | 0.464 | 0.555 | 7.768 | 6.936 | 6.659 | 6.919 | 216.727 | 121.097 | 100.373 | 119.565 |
| Lgals3 | 0.558 | 0.336 | 0.476 | 6.016 | 5.175 | 4.442 | 4.945 | 63.647 | 35.029 | 20.815 | 29.692 |
| Eif4e2 | 0.558 | 0.376 | 0.470 | 8.059 | 7.218 | 6.649 | 6.971 | 265.480 | 147.491 | 99.685 | 123.949 |
| Psat1 | 0.558 | 0.562 | 0.547 | 7.511 | 6.669 | 6.679 | 6.641 | 181.252 | 100.509 | 101.852 | 98.438 |
| Hist1h4h | 0.557 | 0.390 | 0.508 | 4.587 | 3.743 | 3.228 | 3.609 | 23.006 | 12.354 | 8.405 | 11.155 |
| Sars | 0.550 | 0.485 | 0.526 | 6.941 | 6.078 | 5.899 | 6.015 | 121.788 | 66.355 | 58.860 | 63.422 |
| Use1 | 0.549 | 0.372 | 0.468 | 6.902 | 6.036 | 5.476 | 5.806 | 118.538 | 64.460 | 43.680 | 54.724 |
| Rpp25 | 0.547 | 0.406 | 0.433 | 7.000 | 6.129 | 5.699 | 5.791 | 126.907 | 68.787 | 51.115 | 54.189 |
| Tmem238 | 0.544 | 0.442 | 0.562 | 4.048 | 3.169 | 2.869 | 3.217 | 15.526 | 7.971 | 6.328 | 8.267 |
| Mvd | 0.544 | 0.456 | 0.469 | 5.674 | 4.795 | 4.542 | 4.581 | 50.005 | 26.680 | 22.379 | 22.840 |
| Cnpy2 | 0.540 | 0.483 | 0.570 | 7.106 | 6.217 | 6.055 | 6.293 | 136.604 | 73.191 | 65.695 | 77.162 |
| Ptpmt1 | 0.530 | 0.585 | 0.565 | 4.814 | 3.898 | 4.040 | 3.991 | 27.106 | 13.865 | 15.501 | 14.843 |
| Yars | 0.524 | 0.593 | 0.629 | 6.896 | 5.963 | 6.142 | 6.227 | 118.022 | 61.224 | 69.865 | 73.665 |
| Rdm1 | 0.521 | 0.381 | 0.464 | 6.767 | 5.826 | 5.375 | 5.660 | 107.839 | 55.570 | 40.635 | 49.371 |
| Ldlr | 0.516 | 0.527 | 0.556 | 5.542 | 4.587 | 4.619 | 4.696 | 45.549 | 22.962 | 23.649 | 24.814 |
| Gcat | 0.508 | 0.410 | 0.474 | 5.811 | 4.835 | 4.525 | 4.733 | 55.091 | 27.459 | 22.103 | 25.491 |
| March9 | 0.501 | 0.463 | 0.562 | 4.201 | 3.203 | 3.092 | 3.370 | 17.388 | 8.186 | 7.555 | 9.304 |
| Pgp | 0.498 | 0.313 | 0.416 | 6.355 | 5.349 | 4.680 | 5.090 | 80.811 | 39.647 | 24.728 | 32.938 |
| Mrps34 | 0.493 | 0.308 | 0.465 | 6.659 | 5.639 | 4.961 | 5.555 | 99.932 | 48.716 | 30.266 | 45.821 |
| Nsdhl | 0.491 | 0.531 | 0.522 | 5.027 | 4.000 | 4.115 | 4.089 | 31.581 | 14.962 | 16.381 | 15.955 |
| Fdps | 0.453 | 0.421 | 0.416 | 8.185 | 7.044 | 6.937 | 6.921 | 289.726 | 130.611 | 121.929 | 119.673 |
| Usf2 | 0.396 | 0.533 | 0.621 | 4.931 | 3.594 | 4.024 | 4.244 | 29.478 | 11.045 | 15.323 | 17.887 |
| Vax2 | 0.343 | 0.272 | 0.323 | 5.501 | 3.959 | 3.626 | 3.871 | 44.264 | 14.512 | 11.388 | 13.581 |
